# CenSegNet: a generalist high-throughput deep learning framework for centrosome phenotyping at spatial and single-cell resolution in heterogeneous tissues

**DOI:** 10.1101/2025.09.15.676250

**Authors:** Jiaoqi Cheng, Keqiang Fan, Miles Bailey, Xin Du, Rajesh Jena, Constantinos Savva, Ewan Reed, Mengyang Gou, Peixin Zuo, Ramsey Cutress, Stephen Beers, Xiaohao Cai, Salah Elias

## Abstract

Centrosome abnormalities (CA) are a hallmark of epithelial cancers, yet their spatial complexity and phenotypic heterogeneity remain poorly resolved due to limitations in conventional image analysis. We present CenSegNet (Centrosome Segmentation Network), a modular deep learning framework for high-resolution, context-aware segmentation of centrosomes and epithelial architecture across diverse tissue types. Integrating a dual-branch architecture with uncertainty-guided refinement, CenSegNet achieves state-of-the-art performance and generalisability across both immunofluorescence and immunohistochemistry modalities, outperforming existing models in accuracy and morphological fidelity. Applied to tissue microarrays (TMAs) containing 911 breast cancer sample cores from 127 patients, CenSegNet enables the first large-scale, spatially resolved quantification of numerical and structural CA at single-cell resolution. These CA subtypes are mechanistically uncoupled, exhibiting distinct spatial distributions, age-dependent dynamics, and associations with histological tumour grade, hormone receptor status, genomic alterations, and nodal involvement. Structural CA levels are additionally associated with overall survival, supporting the clinical relevance of spatially resolved CA patterns. Discordant CA profiles at tumour margins are linked to local aggressiveness and stromal remodelling. To support broad adoption and reproducibility, CenSegNet is released as an open-source Python library. Together, our findings establish CenSegNet as a scalable, generalisable platform for spatially resolved centrosome phenotyping in intact tissues, enabling systematic dissection of the biology of this organelle and its dysregulation in cancer and other epithelial diseases.

## Introduction

Centrosomes, composed of a pair of orthogonally arranged centrioles surrounded by pericentriolar material (PCM), function as the principal microtubule-organising centres (MTOCs) in animal cells. They play essential roles in diverse cellular processes, including vesicular trafficking, cell polarity, motility, ciliogenesis, and the assembly of a bipolar mitotic spindle during cell division^1, 2^. Centrosome number and size are tightly regulated during the cell cycle, with duplication occurring once per cell cycle during S phase, ensuring the formation of the mitotic spindle and equal inheritance of chromosomes by daughter cells^3–5^.

Centrosome abnormalities (CA) can lead to multipolar spindle formation, chromosomal missegregation, and aneuploidy^3, 6, 7^—a hallmark of cancer^3, 4, 7^. The hypothesis that CA-induced aneuploidy contributes to tumorigenesis was first proposed by Theodor Boveri over a century ago^8^. In recent years, CA have been documented in several solid tumours including breast, prostate, colon, ovarian, and pancreatic cancers^3, 7, 9–13^, as well as haematological malignancies such as multiple myeloma, lymphomas, and leukaemias^14, 15^. While its role in tumour initiation remains debated^6, 9, 16–18^, CA are consistently associated with aggressive disease features, including high-grade histology, poor prognosis, recurrence, and metastasis^3, 7, 9, 19^. Despite their clinical relevance, CA remain poorly characterised at scale due to the lack of robust, high-throughput tools capable of resolving centrosome phenotypes in complex tissue architecture.

Mechanistically, CA can arise from both numerical and structural centrosome defects. Numerical amplification results from centriole overduplication, *de novo* centriole assembly, cytokinesis failure, mitotic slippage and cell–cell fusion^20–29^. Disruption of cell-cycle progression, such as prolonged G2 arrest, can trigger premature centriole disengagement and reduplication *via* PLK1 activation^30, 31^. Fragmentation of the PCM, driven by dysregulation of proteins such as pericentrin, γ-tubulin, PLK4, PLK1, and Aurora-A, also contributes to numerical CA^7, 32–34^. Structural CA, on the other hand, involve aberrant accumulation of PCM^4, 7, 9, 35, 36^, centrosome clustering^3, 9^, or defects in centriole architecture^37–39^. Among these centriole architectural defects, over-elongation and fragmentation can lead to unstable centriole structures and further overduplication^37^, suggesting a mechanistic link between numerical and structural centrosome defects. Yet, their differential contributions to cancer biology remain to be determined.

Manual centrosome annotation remains the standard but is time-consuming, low-throughput, and prone to observer bias. Semi-automated pipelines have emerged to address these limitations. For example, CenFind—a deep learning pipeline based on SpotNet—accurately detects and counts centrioles in cultured cells using immunofluorescence images but does not support structural phenotyping^40^. Other machine learning–based approaches have quantified centriole number and linked supernumerary centrioles to PCM expansion in breast cancer cells^41^. Semi-automated frameworks have also been developed for centrosome quantification in human breast histological sections, including a HistoQuest-aided method detecting structural CA^42^ and an IMARIS-based pipeline integrating both structural and numerical CA^7, 43^. Similarly, a 3D imaging-based pipeline quantified structural centriole abnormalities across cancer types^37^. However, these approaches require manual curation and offer only moderate throughput. A recent high-throughput platform using a Harmony^TM^ software-based framework revealed heterogeneous CA phenotypes in ovarian cancer tissues^44^, yet lacked single-cell resolution and subtype discrimination. Moreover, most existing tools are tailored to immunofluorescence imaging and are not compatible with standard chromogenic immunohistochemistry workflows used in clinical pathology, limiting their diagnosis and translational utility.

To address these limitations, we developed CenSegNet (Centrosome Segmentation Network), a versatile deep learning framework for fully automated, pixel-level detection and segmentation of centrosomes in both immunohistochemistry and immunofluorescence images at single-cell resolution. CenSegNet integrates three state-of-the-art models: Ultralytics YOLOv11, a recent evolution of the You Only Look Once family optimised for fast and accurate performance^45^, U-Net, an encoder–decoder convolutional network designed for precise pixel-wise segmentation^46^, and StarDist for shape-aware instance segmentation pipeline that models objects as star-convex polygons to improve instance separation in dense cellular contexts^47^, enabling robust delineation of epithelial cell boundaries within histopathological specimens. This systematically engineered architecture supports multiscale analysis of centrosomal features in morphologically complex tissue environments. We present a publicly accessible, expert-annotated dataset comprising human and murine breast tissues and human mammary epithelial cell cultures (MECs). Implemented in Python 3.10 with a PyQt5-based graphical interface, CenSegNet enables streamlined data input, real-time PyTorch-based inference, and modular extensibility. Benchmarking against expert annotations and alternative models, CenSegNet achieves pathologist-level accuracy across imaging modalities. Using CenSegNet, we perform the first high-throughput, spatially resolved quantification of numerical and structural CA in clinical breast carcinomas. Our analyses reveal that these CA subtypes are mechanistically uncoupled and evolve along orthogonal spatial gradients: numerical CA predominate in proliferative tumour cores, whereas structural CA accumulate at invasive margins, reflecting distinct evolutionary pressures and microenvironmental cues. These spatial trajectories correlate with histological grade, hormone receptor status, HER2 expression, nodal involvement, and germline alterations, underscoring the role of centrosome dysregulation in driving intratumoral heterogeneity and progression. Notably, variation in structural CA levels is associated with overall survival, highlighting the potential clinical relevance of spatially resolved centrosome phenotyping. Importantly, we validated CenSegNet in other human epithelial tissues including kidney, colon, and appendix, demonstrating its generalisability and potential for broad application in spatial pathology and organelle-level phenotyping across diverse healthy and disease tissue contexts. To support widespread adoption, CenSegNet is released as an open-source Python library, available at https://github.com/SKELab/CenSegNet/ and https://zenodo.org/records/17131573.

## Results

### Development of CenSegNet for robust centrosome segmentation across imaging modalities

We generated tissue microarrays (TMAs) comprising 911 breast tissue cores from normal breast tissue, breast tumours, and adjacent areas, sampled from 127 patients enrolled in the ethically approved and clinically well-characterised BeGIN cohort (Investigating Outcomes from Breast Cancer: Correlating Genetic, Immunological and Nutritional Predictors), from University Hospital Southampton (UHS) (**Fig. 1a**; see Methods). Immunohistochemistry was performed using pericentrin antibody to label centrosomes’ PCM, with haematoxylin counterstaining for nuclei (**Fig. 1a**). For training dataset construction, we manually annotated 14,679 centrosomes within 2,486 epithelial and 108 stromal compartments across 108 selected images (**Fig. 1a**, see Methods). Despite extensive optimisation, co-labelling of TMAs for the centriolar protein GT335 failed to produce reliable or specific signal. This limitation highlights well-recognised challenges associated with detecting centriolar markers in paraffin-embedded tissue sections using chromogenic immunohistochemistry. To address this constraint, we complemented the TMA-based immunohistochemistry workflow with immunofluorescence coupled to high-resolution imaging. These analyses were performed on human MECs using confocal and Lightning-based computational super-resolution imaging on the Leica STELLARIS platform (see Methods), and on mouse mammary epithelium using confocal imaging, with pericentrin and GT335 used to label centrosomes and centrioles, respectively, and DAPI employed as a nuclear counterstain. From this, an immunofluorescence training dataset was assembled comprising 1,285 annotated centrosomes from 143 cells from mouse tissue and 841 human MECs, revealing strong segmentation concordance between pericentrin and GT335 labelling [human MECs: R^2^ = 0.9666 (MCF10A), 0.9085 (MCF10A-PLK4), 0.9995 (MCF10DCIS); mouse tissue: R^2^ = 0.9954] (**Fig. 1b, Supplementary Fig. 1a–h**). We did not detect acentriolar MOTCs that are only labelled for pericentrin. Importantly, as detailed in the Methods section, were further independently validated by two annotators with complementary expertise in cell biology (annotator 1) and computational analysis (annotator 2) in subset of datasets. This validation demonstrated high intra- and inter-observer consistency across datasets. In the immunohistochemistry datasets, mean F1 scores and Cohen’s κ were close to or exceeded 0.90, with a mean IoU of approximately 0.70 (**Supplementary Fig. 2a, b**). In the immunofluorescence datasets, mean F1 scores and Cohen’s κ were close to or exceeded 0.95, with a mean IoU of approximately 0.92 (**Supplementary Fig. 2c, d**). For epithelial segmentation, mean F1 scores and Cohen’s κ were close to or exceeded 0.91, with a mean IoU of approximately 0.87 (**Supplementary Fig. 2e, f**).

**Fig. 1.**
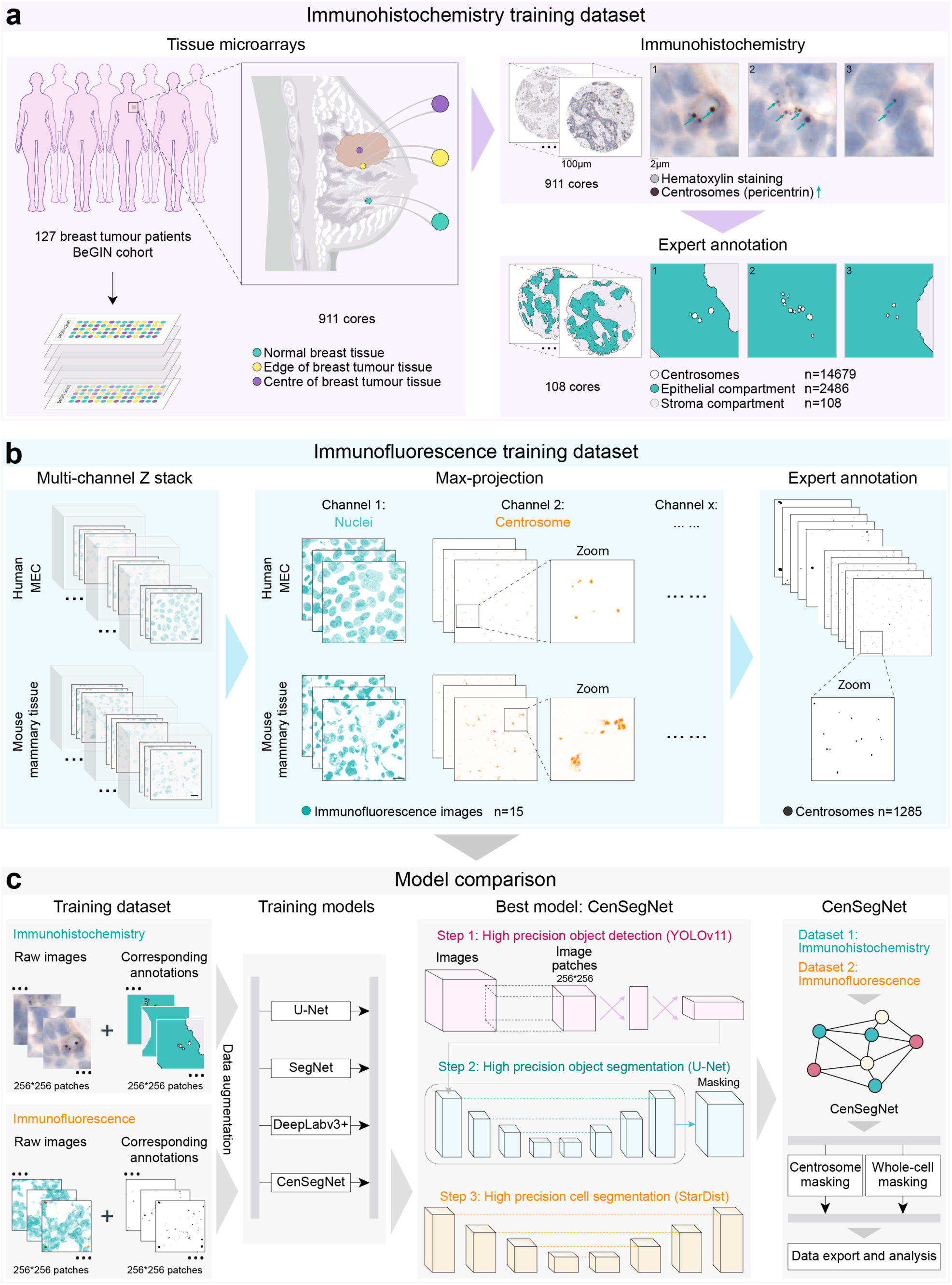
Development and benchmarking of CenSegNet for centrosome segmentation. **a** Left: workflow for generating tissue microarrays (TMAs) comprising 911 breast tissue sample cores from normal breast tissue, breast tumours, and adjacent non-tumour tissue, collected from 127 breast cancer patients in the BeGIN cohort. Figure 1a is schematic for illustrative purposes to demonstrate sampling strategy for the TMA. Cores were taken from FFPE blocks following breast cancer surgery. Top right: immunohistochemical staining of all ROIs with pericentrin (centrosome marker) and haematoxylin (nuclear counterstain). Bottom right: random selection of 108 ROIs used to construct a training dataset containing 2,486 epithelial compartments, 108 stromal compartments and 14,679 annotated centrosomes. **b** Left and middle: representative confocal images of human mammary epithelial cells (MECs) and mouse mammary tumour tissues exhibiting normal or amplified centrosomes. Cells and tissues were stained for pericentrin (orange) and counterstained with DAPI (DNA, teal). Scale bars, 10 µm. Right: training dataset derived from these images, comprising 1,285 annotated centrosomes. **c** Left: 256 × 256-pixel cropped patches from immunohistochemistry and immunofluorescence datasets. Middle left: training of existing segmentation architectures (U-Net, SegNet, DeepLabv3+) and the proposed CenSegNet using these datasets. Middle right: CenSegNet performance across both immunohistochemistry and immunofluorescence test sets. CenSegNet operates in three sequential phases: (1) object detection to generate bounding-box predictions for individual centrosomes; (2) pixel-level segmentation of detected objects; (3) StarDist-based whole-cell segmentation.

Using these datasets, we initially benchmarked established segmentation models. To ensure the models receive appropriately normalised inputs for immunohistochemistry and immunofluorescence imaging modalities—thereby improving generalisability and minimising modality-driven bias—we adopted a modality-specific calibration strategy (see Methods). U-Net, an encoder–decoder convolutional neural network optimised for pixel-wise segmentation^46^, was selected for its extensive use in biomedical imaging. In our datasets, U-Net detected 82.98% of centrosomes in immunohistochemistry images and 97.6% in immunofluorescence, but despite achieving an overall F1 score of 0.85 in immunofluorescence, often the model either under-predicted or over-predicted centrosomes (**Supplementary Fig. 3a, b**). We next evaluated SegNet (**Fig. 1c**), another encoder-decoder model leveraging max-pooling indices for efficient upsampling^48^. It achieved 72.73% and 85% detection in immunohistochemistry and immunofluorescence, respectively, with a precision of 0.90 but reduced recall (0.75) in immunofluorescence and a suboptimal overall F1 score of 0.68 in immunohistochemistry (0.82 in immunofluorescence) (**Supplementary Fig. 3c, d**). We also evaluated DeepLabv3+, a semantic segmentation model utilising atrous convolutions and atrous spatial pyramid pooling for multi-scale contextual information^49^. Despite its proven success in complex image domains, DeepLabv3+ achieved only 63.25% and 75.1% detection rates in immunohistochemistry and immunofluorescence, respectively (**Supplementary Fig. 3e, f**). These findings highlight the limitations of conventional segmentation pipelines when applied to structurally heterogeneous tissues.

To overcome these constraints, we developed CenSegNet, a novel modular framework that integrates object robust detection and segmentation (**Fig. 1c**). Rather than relying on whole-slide segmentation, CenSegNet focuses on single-cell regions while modelling the entire ROI, explicitly integrating object detection with region-based segmentation in a three-step architecture (**Fig. 1c**). This design enables a more comprehensive capture of centrosome spatial distributions and morphological features, addressing key challenges such as occlusion, small object detection, and the disambiguation of overlapping structures. The framework, first employs YOLOv11^45^, a state-of-the-art object detection model, to identify potential centrosome candidates. YOLOv11 comprises a convolutional backbone for feature extraction, a neck for multi-scale aggregation, and a head for classification and localisation. We fine-tuned YOLOv11-seg^45, 50^, a segmentation-optimised variant to further enhance detection accuracy, and applied a range of data augmentations—including hue, saturation, and value (HSV) adjustments, as well as geometric transformations such as translation, scaling, shearing, and horizontal flipping—to enhance generalisability (**Fig. 1c**; see Methods). To refine segmentation, we integrated U-Net in the second step. For all detected centrosomes, we extracted centrosome-centred patches (256 × 256 pixels, with 40-pixel padding) as input to a U-Net skip-connected encoder-decoder architecture for precise delineation. These patches underwent similar augmentation strategies as in the detection step, ensuring consistency. To accurately quantify centrosome numbers per cell, we incorporated StarDist^47^ in the third step, a deep learning based instance segmentation tool widely adopted for nuclear and cell boundary segmentation in biomedical imaging (**Fig. 1c**). In our immunohistochemistry datasets, StarDist segmented 764,354 cells (**Supplementary Fig. 1i**). To validate StarDist-aided cell-level centrosome assignment in complex tissues, we analysed keratin 8 (KRT8)-labelled mouse immunofluorescence (620 cells) and human immunohistochemistry (620 cells) tissues. StarDist-based segmentation demonstrated strong concordance with KRT8 (immunofluorescence: R^2^ = 0.99; immunohistochemistry: R^2^ = 0.9005) (see **Supplementary Fig. 1g-k**).

Collectively, these results demonstrate that CenSegNet’s multistep architecture effectively overcomes the limitations of conventional segmentation approaches, delivering robust, generalisable centrosome detection and quantification across diverse imaging modalities and complex tissue contexts.

### Validation of CenSegNet for scalable and high-precision centrosome segmentation

To evaluate CenSegNet’s performance, we compiled independent test datasets of 25 immunohistochemistry TMA cores and 17 immunofluorescence images. Predicted centrosome counts were compared to manually annotated ground truth, revealing strong correlations in both immunohistochemistry (R^2^ = 0.9999) and immunofluorescence (R^2^ = 0.9873) (**Fig. 2a–c**). CenSegNet consistently outperformed U-Net, SegNet, and DeepLabv3+ in segmentation accuracy and boundary resolution across both modalities, with the YOLOv11 detection module significantly enhancing overall performance (**Fig. 2a–c**). To further assess precision, we used an independently annotated subset of 550 centrosomes from tumour regions in the immunohistochemistry test set. Again, CenSegNet demonstrated superior pixel-level segmentation compared to all benchmarks (**Fig. 2d**). F1 score analysis across modalities yielded a mean of 0.82 for CenSegNet, outperforming U-Net (0.72), SegNet (0.68), and DeepLabv3+ (0.65) (**Fig. 2a–e**). We next benchmarked performance on 6,475 expert-annotated centrosomes—921 from normal tissue, 2,694 from edge regions, and 2,860 from tumour cores (**Fig. 2f**). These annotations, stratified by size (0.5–1.0 µm^2^ to >10.5 µm^2^), were compared with the full cohort of 333,148 automatically segmented centrosomes (0–0.5 µm^2^ to >12.0 µm^2^)—10,834 from normal tissue, 91,803 from edge tissue, and 230,511 from tumour tissue (**Fig. 2g**). Both datasets revealed similar size distributions, with the majority falling within 2.0–3.0 µm^2^ and a peak at 2.0–2.5 µm^2^. Normal tissue showed a higher proportion of 1.0–2.0 µm^2^ centrosomes, while edge regions had more large centrosomes than either normal or tumour cores. Using a previously established threshold of 6.5 µm^2^-size to define structural CA in human breast cancer tissues^42^ (see Methods), both datasets confirmed that centrosomes exceeding this size were absent in normal tissue.

**Fig. 2.**
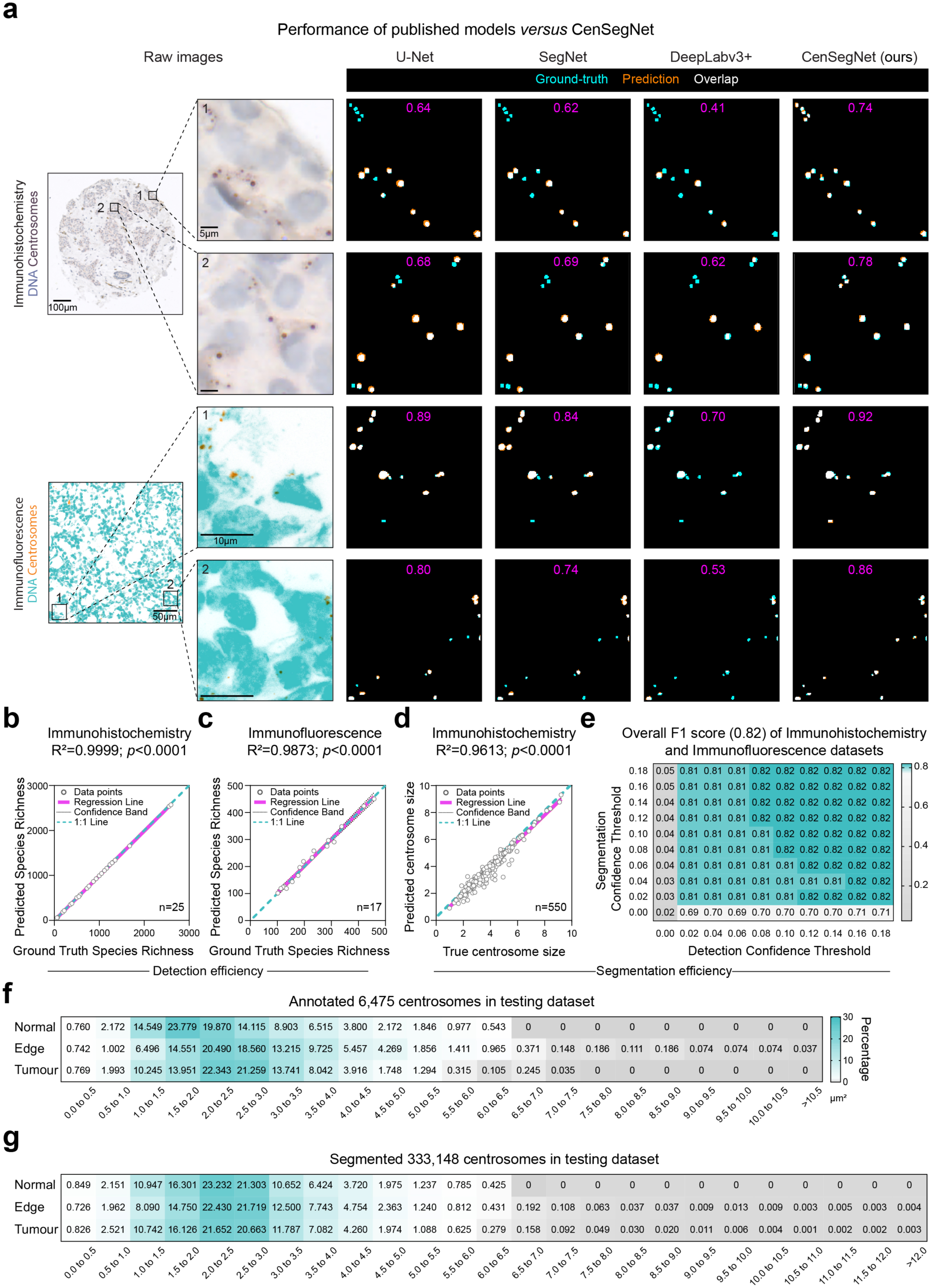
Validation of CenSegNet performance against alternative models and human annotations. **a** Left: representative immunohistochemistry and immunofluorescence images used for centrosome segmentation. Right: colour overlays showing predictions from U-Net, SegNet, DeepLabv3+, and CenSegNet (teal: ground-truth annotations; orange: predicted segmentation; white: overlap between predictions and ground-truth). Magenta text indicates the F1 score for corresponding images of each method (0: complete disagreement; 1: complete concordance). Yellow arrow marks the overprediction. **b** Pearson correlation between centrosome counts predicted by CenSegNet and ground truth across 25 immunohistochemistry images. Grey circles represent individual patients, purple line represents regression line, Black dot line represents confidence band, and teal dot line represents 1:1 line. Two-tailed Pearson correlation test followed by simple linear regression for visualisation, \*\*\*\**P* < 0.0001. **c** Pearson correlation between centrosome counts predicted by CenSegNet and ground truth across 17 immunofluorescence images. Two-tailed Pearson correlation test, followed by simple linear regression for graphical representation, \*\*\*\**P* < 0.0001. **d** Pearson correlation between centrosome size predicted by CenSegNet and measured by annotators (n = 550 centrosomes from >200 ROIs across 20 images). Two-tailed Pearson correlation test, followed by simple linear regression for graphical representation, \*\*\*\**P* < 0.0001. **e** Heatmap of overall F1 scores (maximum 0.82) from combined immunohistochemistry and immunofluorescence test datasets. The *x*-axis represents the detection confidence threshold, and the *y*-axis represents the segmentation confidence threshold. Each cell represents the mean F1 score for that threshold pair; darker cyan indicates higher values. Optimal performance (F1 = 0.82) was achieved across a broad range of threshold combinations, indicating model robustness to parameter variation. **f, g** Manual annotations of 6,475 centrosomes from normal, edge, and tumour regions—stratified by size (0.5–1.0 μm^2^ to >10.5 μm^2^) **(f)** were compared with automated segmentations from the full dataset of 333,148 centrosomes **(g)** (0–0.5 μm^2^ to >12.0 μm^2^). CenSegNet segmentation achieved performance comparable to expert human annotation. Data are presented as individual data points. Source data are provided as Source Data file.

We further assessed the sensitivity of segmentation performance to the choice of detection and segmentation thresholds across four representative image types: immunohistochemistry, confocal immunofluorescence, lightning-based super-resolution immunofluorescence, and haematoxylin-stained images. For each image type, either the detection or segmentation threshold was varied from 0.1 to 0.8 while the other threshold was fixed at 0.15 (default value), and performance was quantified using F1 score, IoU and Cohen’s κ (**Fig 3**). Across all modalities, CenSegNet exhibited robust performance over varying threshold values. In immunohistochemistry images, performance remained stable as the detection threshold increased from 0.1 to 0.5, with reductions observed only at higher thresholds (**Fig. 3a, b**). When the segmentation threshold was varied, F1 score, IoU, and Cohen’s κ remained stable across the tested range (**Fig. 3a, c**). A similar pattern was observed for both confocal (**Fig. 3d-f**) and lightning-based super-resolution (**Fig. 3g-i**) immunofluorescence images. F1 score and Cohen’s κ remained stable across detection thresholds from 0.1 to 0.8, with only minor fluctuations in IoU (**Fig. 3d, e, g, h**), while variation of the segmentation threshold over the same range had little effect on performance (**Fig. 3d, f, g, i**), indicating that segmentation accuracy was not sensitive to threshold selection. For epithelial segmentation in haematoxylin-stained images, performance was stable across detection thresholds from 0.1 to 0.3 and decreased at higher values (**Fig. 3j, k**). Similarly, segmentation performance remained stable when the segmentation threshold was varied from 0.1 to 0.4, with decreases observed only at larger values (**Fig. 3j, l**). Notably, the default threshold of 0.15 consistently fell within the stable performance range across all evaluated image types and metrics. Together, these validation analyses establish CenSegNet as a scalable, high-precision framework for centrosome segmentation that delivers robust and consistent performance across imaging modalities, tissue compartments, and size distributions, supporting its applicability to high-throughput analysis in complex tissue environments.

**Fig. 3.**
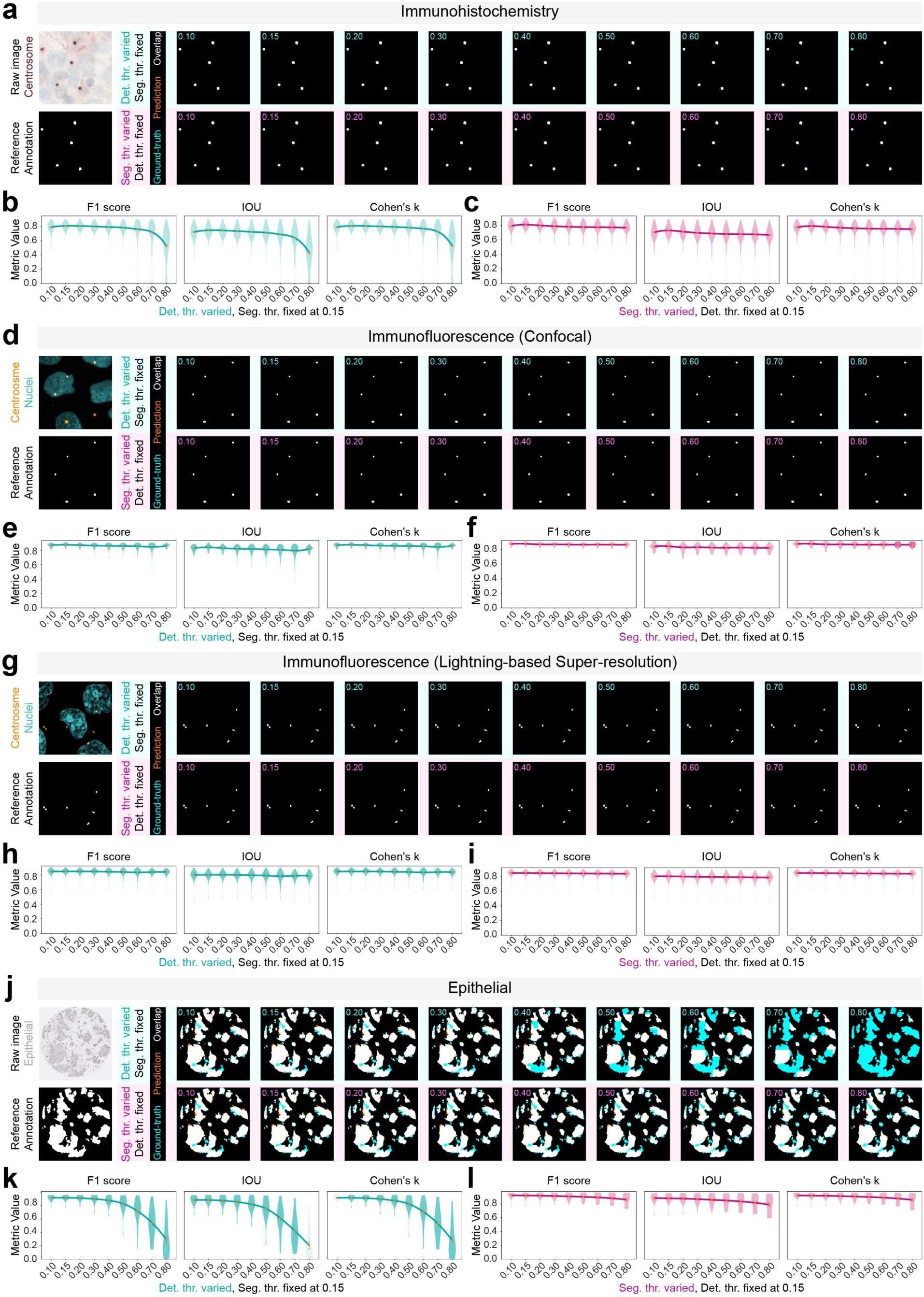
Effect of detection and segmentation thresholds on centrosome and epithelial segmentation performance of CenSegNet. **a, d, g, j** Representative images and corresponding reference annotations shown alongside CenSegNet predictions generated using different threshold settings. **a** Immunohistochemistry image of centrosome staining in human breast tumour tissue. **d, g** Representative confocal and lightning-based super-resolution immunofluorescence images of centrosome staining. **j** Representative haematoxylin-stained (epithelial) image of human breast tumour tissue used for epithelial region segmentation. In the upper row of each panel, the detection threshold (Det. thr) was varied while the segmentation threshold (Seg. thr) was fixed at 0.15. In the lower row, the segmentation threshold was varied while the detection threshold was fixed at 0.15. Cyan indicates the reference annotation; orange indicates the prediction and white indicates overlap. **b, e, h, k** F1 score, intersection over union (IoU) and Cohen’s κ across varying detection thresholds with the segmentation threshold fixed at 0.15. **c, f, i, l** F1 score, IoU and Cohen’s κ across varying segmentation thresholds, with the detection threshold fixed at 0.15. Violin plots show the distribution of metric values across images; lines indicate the mean trend across thresholds. For **b, c** data were annotated from 200 256 × 256-pixel patches from 25 immunohistochemistry images. For **e, f** data were annotated from 60 1,024 × 1,024-pixel patches from 10 confocal immunofluorescence images. For **h, i** data were annotated from 60 1,024 × 1,024-pixel patches from 10 lightning-based super-resolution immunofluorescence images. For **k, l** data were annotated from 19 images ranging from 4,440 × 4,492 to 6,593 × 7,360 pixels. Source data are provided as a Source Data file.

### CenSegNet reveals orthogonal and spatially organised numerical and structural CA in breast cancer

Our robust orthogonal validation above demonstrating that CenSegNet segments bona fide centrosomes provides a grounded rationale for high-throughput quantitative analyses distinguishing between numerical CA (Num CA; increased number of centrosomes per cell) and structural CA (Stru CA; altered size and/or organisation of the PCM) based on pericentrin labelling in our immunohistochemistry TMA data, to dissect their relative contributions in breast cancer. Num CA and Stru CA were not correlated within either tumour edge or core regions (**Supplementary Fig. 4a**), suggesting mechanistic independence. At the single-cell level, increasing centrosome number per cell was associated with a reduction in the size of individual centrosomes (**Supplementary Fig. 4b**), which become smaller, more uniform in cells containing >4 centrosomes—implying a compensatory constraint on total centrosome volume. These observations suggest that Stru CA and Num CA represent orthogonal axes of centrosome dysregulation. In contrast, Num CA levels correlated positively between edge and tumour compartments (R^2^ = 0.4857; **Supplementary Fig. 4c**), indicating progressive numerical amplification from tumour margins inward. This was reflected in an increasing proportion of cells with ≥4 centrosomes from normal tissue to the edge and tumour core (**Fig. 4a**). Stratifying patients into four CA phenotypes (Stru⁻Num⁻, Stru⁺Num⁺, Stru⁺Num⁻, Stru⁻Num⁺) revealed that CA was widespread, detected in 89.8% of edge and 95% of tumour regions (**Supplementary Fig. 4d**). Notably, 73% of tumour regions exhibited both Stru CA and Num CA, representing a ∼19.3% increase compared to edge regions (**Supplementary Fig. 4d**). While the Stru⁺Num⁻ group accounted for 18.4% of edge regions but only 2% of tumour regions, the opposite trend was observed for the Stru⁻Num⁺ group (10.2% *versus* 20%, respectively). These findings allow us to conclude that structural and numerical CA are mechanistically uncoupled and highlight a spatial shift from centrosome enlargement at the tumour edge to centrosome accumulation in the tumour core.

**Fig. 4.**
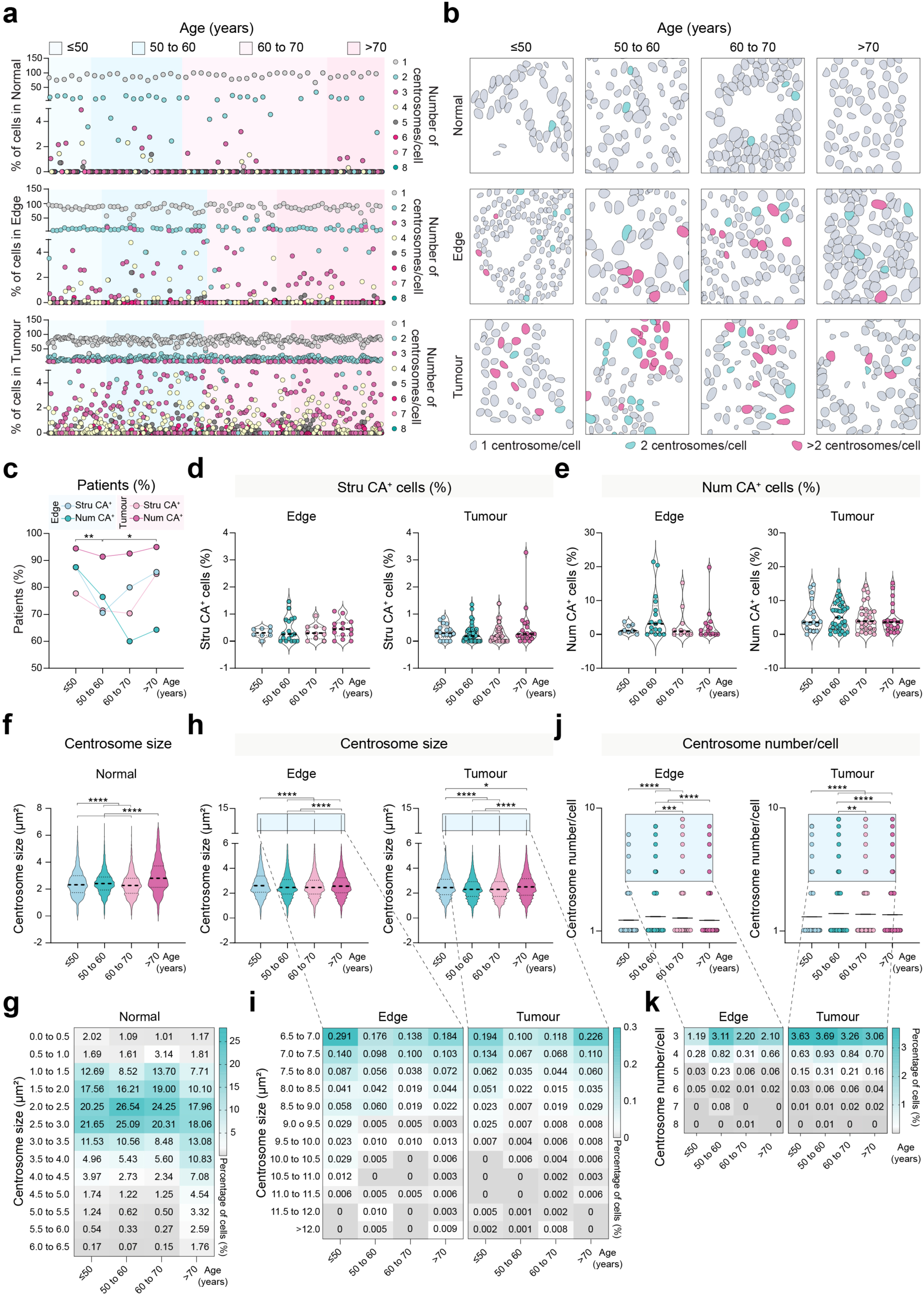
Age-related gradients of centrosome abnormalities across breast tissue regions. **a** Percentage of cells with different centrosome numbers in normal, edge, and tumour regions. **b** Representative cell segmentation masks of normal, edge, and tumour regions across patient age groups (Light grey: cells with one centrosome; purple: cells with two centrosomes; cyan: cells with more than two centrosomes). **c** Percentage of patients with structural CA (Stru CA) and Num CA across age groups in edge (≤50 years: n = 8 patients; 50–60 years: n = 17 patients; 60–70 years: n = 10 patients; >70, n = 14 patients) and Tumour (≤50 years: n = 18 patients; 50–60 years: n = 35 patients; 60–70 years: n = 27 patients; >70 years: n = 20 patients) regions. Two-way ANOVA with Tukey’s test, *\*P* = 0.0477, \*\**P* = 0.0053. Data are presented as mean ± s.e.m. **d** Percentage of cells with Stru CA in edge and tumour regions across age in Edge and Tumour regions. One-way ANOVA (Stru CA^+^ cells: Edge, *P* = 0.7637; Tumour, *P* = 0.4489 with Tukey’s test. absence of asterisks indicates no statistically significant correlation. **e** Percentage of cells with Num CA in edge and tumour regions across age. One-way ANOVA (Num CA^+^ cells: Edge, *P* = 0.3427; Tumour, *P* = 0.9086) with Tukey’s test, absence of asterisks indicates no statistical significance. Data are presented as mean ± s.e.m. **f, g** Centrosome segmentation results from normal regions across age groups, stratified by size (0.5–1.0 μm^2^ to 6.0–6.5 μm^2^). One-way ANOVA (\*\*\*\**P* < 0.0001) with Tukey’s test, \*\*\*\**P* < 0.0001. Data are presented as violin plots showing the distribution of values; dashed lines indicate median and interquartile ranges. **h, i** Centrosome segmentation results from edge and tumour regions across age groups, stratified by size (6.5–7.0 μm^2^ to 12.0 μm^2^). One-way ANOVA (\*\*\*\**P* < 0.0001) with Tukey’s test, Edge: \*\*\*\**P* < 0.0001; Tumour: \**P* = 0.0484, \*\*\*\**P* < 0.0001. Data are presented as violin plots showing the distribution of values; dashed lines indicate median and interquartile ranges. **j, k** Centrosome number per cell in edge and tumour regions across age groups, stratified by centrosome number (1–8). One-way ANOVA (Edge, \*\*\*\**P* < 0.0001; Tumour, \*\*\*\**P* < 0.0001) with Tukey’s test, Edge: \*\*\**P* = 0.0006, \*\*\*\**P* < 0.0001; Tumour: \*\**P* = 0.0054, \*\*\*\**P* < 0.0001, absence of asterisks indicates no statistical significance. Data are presented as individual data points and mean ± s.e.m. Source data are provided as Source Data file.

To directly test the relationship between centriole number, centrosome number, and centrosome size, we performed functional perturbation experiments targeting PLK4, the master regulator of centriole biogenesis^9^. Using a well-established Tet-On MCF10A-PLK4 system, centriole amplification was first induced by doxycycline treatment, followed by pharmacological inhibition of PLK4 activity using the selective inhibitor centrinone^51^. Both parental MCF10A and MCF10A-PLK4 cells were co-labelled for pericentrin (PCM) and GT335 (centriole) and imaged by lightning-based super-resolution imaging to enable simultaneous quantification of centrosome number, centriole number, and centrosome size (**Supplementary Fig. 5a**). Under these conditions, centrinone treatment did not significantly alter the number of centrosomes per cell, as defined by pericentrin-positive foci, in either cell line (**Supplementary Fig. 5b**). In contrast, PLK4 inhibition resulted in a marked reduction in centriole number per centrosome in both MCF10A and MCF10A-PLK4 cells (**Supplementary Fig. 5c**), confirming effective inhibition of PLK4-dependent centriole duplication. Notably, centrinone treatment was also associated with a significant reduction in centrosome area in both contexts (**Supplementary Fig. 5d**), indicating that centrosome size is sensitive to PLK4 activity and centriole content independently of centrosome number. Correlation analyses further supported this dissociation. Under control (DMSO-treated) conditions (**Supplementary Fig. 5e, f**), centrosome area exhibited a strong positive correlation with centriole number per centrosome in both MCF10A (R^2^ = 0.4442) and MCF10A-PLK4 cells (R^2^ = 0.7752) (**Supplementary Fig. 5i, j**), consistent with coordinated scaling between centriole content and PCM expansion. Upon centrinone-mediated PLK4 inhibition (**Supplementary Fig. 5g, h**), this coupling was substantially weakened in both cell types (MCF10A: R^2^ = 0.1941; MCF10A-PLK4: R^2^ = 0.1825; **Supplementary Fig. 5k, l**). Similarly, while centrosome number per cell correlated positively with total centriole number under control conditions (MCF10A: R^2^ = 0.3528; MCF10A-PLK4: R^2^ = 0.3544; **Supplementary Fig. 5m, n**), this relationship was largely lost following centrinone treatment (MCF10A: R^2^ = 0.0405; MCF10A-PLK4: R^2^ = 0.0952; **Supplementary Fig. 5o, p**). Together, these functional perturbation experiments show that PLK4 inhibition disrupts the quantitative relationships linking centriole number, centrosome size, and centrosome number, while centriole content and centrosome size remain tightly coupled under unperturbed conditions. These experiments therefore further support the mechanistic uncoupling of numerical and structural CA observed in breast cancer tissues.

These findings prompted us to consider whether enlarged centrosomal structures might reflect centrosome clustering, a process essential for the survival of cancer cells by which extra centrosomes assemble to form the two poles of the mitotic spindle^3, 9^. Consistent with this, centrosome clustering in our immunohistochemistry data would be expected to manifest as large, irregular pericentrin-labelled PCM regions with internal intensity heterogeneity, indicative of multiple centrosomes in close proximity^52, 53^. Instead, our morphometric analyses show that circularity and solidity were largely maintained across the range of centrosome sizes. Circularity and solidity did not correlate significantly with centrosome size (**Supplementary Fig. 6a**), and enlarged centrosomes did not differ significantly from smaller centrosomes in these parameters (**Supplementary Fig. 6b**). These findings indicate that the majority of enlarged centrosomal foci observed in our tissue samples retain the compact, rounded morphology expected of single centrosomes. While we cannot entirely exclude the possibility that a minority of enlarged structures arise from centrosome clustering, our data suggest that clustering is unlikely to be the predominant explanation for the enlarged centrosomes observed.

### CenSegNet enables spatial profiling of CA subtypes and clinical correlates in breast cancer

To investigate how CA correlates with ages, we stratified patients into four groups: <50, 50–60, 60–70, >70 years. Num CA was elevated in patients aged 50–70 years across edge and tumour regions (**Fig. 4a, b**). The frequency of patients with detectable Num CA and Stru CA at the tumour edge markedly decreased in the 50–70 age group, resurging in patients over 70 years (**Fig. 4c**). Although fewer 50 to 60-year-old patients exhibited detectable CA at the edge, those that did had high levels of both CA subtypes (**Fig. 4d, e**). CA levels in tumour cores were relatively stable across age. Notably, normal tissues exhibited age-dependent increases in centrosome size, especially in patients over 70 (**Fig. 4f, g**), consistent with previous reports linking age to centrosome expansion *via* cumulative DNA damage^54^. In edge and tumour regions, centrosome size decreased between ages 50–70 before rising again in patients over 70 (**Fig. 4h, i**), while centrosome numbers followed the opposite trend, increasing in the 50–70 group before decreasing in the oldest cohort (**Fig. 4j**). This dynamic was mirrored by a higher proportion of cells with three centrosomes or more in the 50–60 age group (**Fig. 4k**). Finally, we did not observe significant differences in age distribution across the four CA-defined patient groups (Stru^−^Num^−^, Stru^+^Num^+^, Stru^+^Num^−^, and Stru^−^Num^+^) (**Supplementary Fig. 7a**). Thus, while age alone may not initiate CA in breast cancer, it modulates its spatial distribution and severity—particularly by promoting numerical amplification in the 50–70 age group and structural amplification in patients over 70.

We next evaluated associations between CA subtypes and clinicopathological features. Tumours lacking CA (Stru⁻Num⁻) displayed smaller total and invasive areas in the tumour core relative to the edge (**Supplementary Fig. 7b, c**), a pattern absent in other CA groups. Structural and numerical CA prevalence increased with histological grade and nodal involvement (**Fig. 5a, b**). Stru CA was inversely associated with tumour size across tumour (T) stages both edge and tumour regions, while Num CA displayed region-specific variation (**Fig. 5c**). Stru CA and Num CA were more frequent at the edge in invasive ductal carcinoma, but more abundant in the core of mixed-subtype tumours (**Fig. 5d**). To further assess CA heterogeneity, patients were classified by composite CA defects into Stru^high^Num^high^, Stru^high^Num^low^, Stru^low^Num^low^, Stru^low^Num^high^ CA groups (**Fig. 5e, f**). CA burden was not age-associated (**Fig. 5g**), and distinct CA subtypes were linked to tumour behaviour. At the edge, Stru^high^Num^low^ and Stru^low^Num^high^ CA tumours were associated with greater nodal involvement than either the Stru^high^Num^low^ or Stru^low^Num^high^ CA groups (**Fig. 5h**), whereas no such differences were observed across CA groups in the tumour core. Stru^low^Num^high^ CA tumours exhibited the smallest overall size, while the largest tumours were enriched in Stru^high^Num^low^ (edge) and Stru^high^Num^high^ (tumour) CA groups (**Fig. 5i, j**). Stru^high^Num^low^ CA tumours grew more aggressively at the edge, while Stru^high^Num^high^ CA tumours expanded predominantly in the core (**Fig. 5k**). While Stru^low^Num^low^ CA was the most frequent subtype among large tumours across both regions, neither Num CA (in tumour cores) nor Stru CA (at the edge) alone stratified tumour size (**Fig. 5l**). About 40% of tumours within the Stru^high^Num^high^ CA group exhibited nodal involvement greater than N1 in the core (**Fig. 5m**), consistent with increased metastatic potential. Many tumours with Stru^low^Num^low^ CA status at the edge required nodal clearance, suggesting local aggressiveness independently of global CA burden (**Fig. 5n**). Identified germline variants were exclusively observed in the Stru^high^Num^high^ CA group (**Fig. 5o**), consistent with associations between CA and BRCA1-driven genomic instability^55^.

**Fig. 5.**
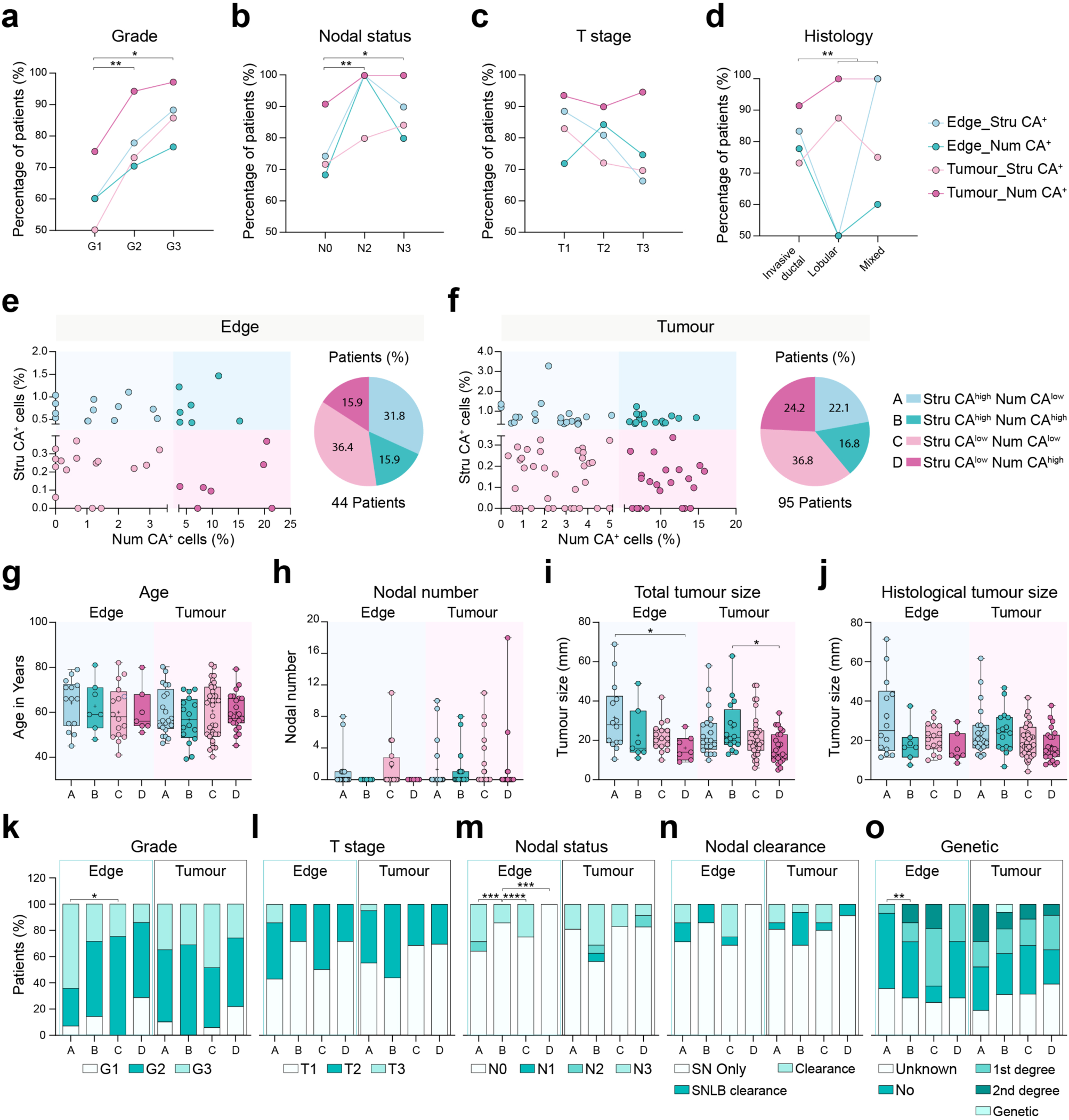
Patient stratification by CA burden and correlation with clinicopathological features. **a–d** Percentage of patients with Stru CA and Num CA across histological tumour grade, nodal status, histological tumour size, and histological tumour type in Edge [Grade: G1 (n= 5), G2 (n=27), G3 (n=17); Nodal status: N0 (n=38), N2 (n=1), N3 (n=10); T stage: T1 (n=18), T2 (n=16), T3 (n=15); Histological type: Invasive ductal (n=36), Lobular (n=8), Mixed (n=5)], and Tumour [Grade: G1 (n=12), G2 (n=52), G3 (n=35); Nodal status: N0 (n=77), N2 (n=4), N3 (n=18); T stage: T1 (n= 30), T2 (n= 29), T3 (n= 40); Histological type: Invasive ductal (n=82), Lobular (n=8), Mixed (n=8)] regions. Two-way ANOVA (Grade, \*\**P* = 0.0020; Nodal status, ** *P* = 0.0030; T stage, *P* = 0.4024; Histological type, ** *P* = 0.0037) with Tukey’s test, Grade: *\*P* = 0.0130, \*\**P* = 0.0018; Nodal status: \**P* = 0.0113, \*\**P* = 0.0030; Histological type: (Invasive ductal *versus* Lobular) \*\**P* = 0.0063, (Invasive ductal *versus* Mixed) \*\**P* = 0.0061. Data are presented as mean ± s.e.m. **e, f** Patients classified by composite CA burden into Stru^high^Num^low^ (A), Stru^high^Num^high^ (B), Stru^low^Num^low^ (C), and Stru^low^Num^high^ (D) groups in Edge (A: n = 14 patients; B: n = 7 patients; C: n = 16 patients; D: n = 7 patients) and Tumour (A: n = 20 patients; B: n = 16 patients; C: n = 35 patients; D: n = 23 patients) regions. Histograms and pie charts show percentages of patients in each group in edge and tumour regions. **g–j** Comparative analysis of patient characteristics, including age, number of involved nodes, and total and histological tumour size, across composite CA groups A–D in edge and tumour regions. One-way ANOVA Age (Edge, *P* = 0.7719; Tumour, *P* = 0.6469) Nodal number (Edge, *P* = 0.3164; Tumour, *P* = 0.9944); Total tumour size (Edge, \**P* = 0.0491; Tumour, \**P* = 0.0318); Histological tumour size (Edge, \**P* = 0.0403; Tumour, *P* = 0.0732) with Tukey’s test, Total tumour size (Edge, \**P* (A vs D) = 0.0484; Tumour, \**P* (B vs D) = 0.0296 ). Absence of asterisks indicates no statistical significance. Data are presented as individual data points and as box and whiskers plots showing the distribution of values, median and quartiles. **k–o** Percentage of patients with different tumour characteristics (histological tumour grade, T stage, nodal status, nodal clearance, genetic subtype) across composite CA groups A–D in edge and tumour regions. Fisher’s exact test, left: \**P* = 0.0261; middle: \*\*\**P* = 0.0005 (middle left), \*\*\**P* = 0.0006 (middle right) and \*\*\*\**P* < 0.0001; right: \*\**P* = 0.0091. **o** 1st degree, family history with 1^st^ degree relative; 2^nd^ degree, family history with 2^nd^ degree relative. Genetic, pathogenic variant in breast cancer-related gene. Source data are provided as Source Data file.

We also evaluated the relationship between CA subtypes and body composition. Tumours with Stru^low^Num^high^ CA in the core were associated with increased fat-free mass index (FFMI), (**Supplementary Fig. 8a**), while fat mass index (FMI), waist circumference, and weight showed no differences across groups or regions (**Supplementary Fig. 8b-d**). Taller patients more frequently exhibited Stru^low^Num^low^ CA status in the edge regions, though this association was not observed in cores (**Supplementary Fig. 8e**). Collectively, these findings indicate that spatial patterns of CA are linked to tumour clinical parameters. Stru^high^Num^high^ CA tumours define a high-risk subgroup marked by large size, nodal spread, and genomic alterations, while Stru^low^Num^low^ CA tumours—despite low CA burden—can display local aggressiveness.

Single-cell analyses revealed further divergent CA subtype dynamics. At the edge regions, Num CA decreased with increasing tumour size (**Supplementary Fig. 9a**), while Stru CA was enriched in poorly differentiated but smaller tumours (**Supplementary Fig. 9b**). In the core, the proportion of cells harbouring either CA subtype increased with both total and invasive tumour size (**Supplementary Fig. 9c, d**). Num CA showed no significant differences across histological tumour grades, although it was more prevalent in well-differentiated tumours (**Supplementary Fig. 9c**). Stru CA levels were enriched in poorly differentiated tumours (**Supplementary Fig. 9d**). No significant subtype-specific CA differences were observed across histological subtypes (**Supplementary Fig. 9a-d**), but cells from mixed tumours harboured more centrosomes in both compartments (**Supplementary Fig. 10**), highlighting elevated Num CA as linked to increased tumour heterogeneity. Together, these findings indicate a dynamic evolution of CA during tumour progression, with early-stage tumours characterised by numerical defects and advanced tumours accumulating structural abnormalities.

We further explored CA patterns in the context of hormone receptor status, a key clinical determinant in breast cancer^56, 57^. Stru CA levels varied by receptor status in both edge and tumour core compartments (**Supplementary Fig. 11a**). HER2⁻ tumours had the highest Stru CA at the edge, while PR⁻ and ER⁻ tumours had the lowest. In contrast, ER⁻ tumours displayed elevated Stru CA in the core, with HER2⁺ tumours showing the lowest levels. Num CA also showed compartment-specific trends: HER2⁺ tumours exhibited high Num CA at the edge but low levels in the core; ER⁻ tumours showed the inverse pattern (**Supplementary Fig. 11b**). Despite broadly similar spatial trends between Stru CA and Num CA across receptor-defined subtypes, HER2⁺ tumours emerged as an exception (**Supplementary Fig. 11c, d**). HER2^moderate^ (2+) tumours had significantly lower Num CA levels than HER2^high^ (3+) tumours in both compartments (**Supplementary Fig. 11e**), suggesting HER2 dosage impacts centrosome number. HER2⁺ tumours were overrepresented in discordant CA phenotypes—Stru^low^Num^high^ (28.6%) and Stru^high^Num^low^ (35.7%)—at the edge (**Supplementary Fig. 11f**), indicating HER2 may differentially regulate centrosome structure and number depending on spatial context. This distribution was not observed in the tumour core. These observations point to spatially resolved, hormone receptor-specific influence of CA patterns, particularly via HER2 signalling, in a microenvironment-dependent manner.

Finally, we examined whether spatially resolved CA patterns are associated with patient survival outcomes, including overall survival (OS) and recurrence-free survival (RFS). We assessed numerical and structural CA within both the tumour core and the tumour–normal edge region. In the edge region, neither Num CA nor Stru CA showed a statistically significant association with OS or RFS in the current cohort (**Supplementary Fig. 12a–d**), and similarly, no significant associations were observed for most CA measures in the tumour compartment (**Supplementary Fig. 12e, g, h**). Notably, however, lower levels of Stru CA in the tumour core were associated with significantly more favourable OS (**Supplementary Fig. 12f**). Beyond this statistically significant association, several additional consistent trends emerged. Tumours exhibiting lower Num CA together with higher Stru CA in the tumour core displayed comparatively poorer OS trends (**Supplementary Fig. 12e**), although these associations did not reach statistical significance. These patterns align with our clinicopathological analyses, in which Num^high^Stru^low^ tumours displayed a more favourable profile, including the smallest tumour sizes (**Fig. 5i**), while tumours with Stru Num^low^Stru^high^ CA exhibited larger tumour size (**Fig. 5j**) and were enriched for a first- or second-degree family history of breast cancer (**Fig. 5o**). Although some Num^high^Stru^low^ tumours exhibited higher absolute node counts, their pathological nodal stage remained N0 (**Fig. 5m**), supporting a less aggressive overall phenotype. To further evaluate potential prognostic relevance, we performed multivariable Cox proportional hazards regression analyses incorporating established clinicopathological covariates, including age, prior breast cancer, smoking status, genetic status, tumour grade, tumour size, T stage, and nodal status (**Supplementary Fig. 13**). After adjustment, neither Num CA nor Stru CA in the tumour core or edge was significantly associated with OS or RFS. Notably, several well-established prognostic variables^58, 59^ likewise did not reach statistical significance, likely reflecting the limited cohort size and number of outcome events^60, 61^. Inclusion of multiple covariates reduced model stability, as indicated by widened confidence intervals and convergence limitations. To maximise interpretability under these constraints, we performed additional exploratory stratification analyses based on combined numerical and structural CA states across spatial compartments. Although these analyses—apart from the association observed for tumour-core Stru CA—did not reach statistical significance, they revealed consistent outcome trends, with lower structural CA in the tumour core associated with more favourable survival patterns, and higher structural CA combined with lower numerical CA associated with adverse outcomes and unfavourable clinicopathological features (**Supplementary Figs 12, 13**). Given that the BeGIN cohort is a recently established study initiated in 2015 (see Methods), the current analyses remain underpowered for definitive prognostic modelling. Taken together, these results identify structural CA within the tumour core as the only CA feature significantly associated with OS in this cohort, while also indicating that spatially resolved numerical and structural CA patterns align with clinically meaningful outcome trends. Consistent with our broader findings, they suggest that elevated structural CA—particularly in the tumour core—may reflect more aggressive disease behaviour, whereas numerical CA in relative isolation appears better tolerated. Collectively, these analyses reinforce the clinical relevance of the orthogonal and spatial organisation of centrosome abnormalities identified in this study and highlight CA subtypes, together with their spatial context, as biologically distinct contributors to breast cancer heterogeneity that warrant validation in larger cohorts with mature follow-up.

### CenSegNet reveals CA patterns linked to tumour subtype and progression

Using CenSegNet-derived spatial quantifications, we compared dynamic shifts in Stru CA and Num CA between tumour edges and cores. Patients were classified based on whether each CA subtype was more abundant in the tumour core (T>E) or at the edge (E>T). Baseline clinicopathological features were similar between spatial groups (**Supplementary Fig. 14**). Stru CA^T>E^ tumours were associated with lobular histology (15.8%), lower histological tumour grade (26.3% grade 3), and lower HER2^+^ prevalence (15.8%). In contrast, Stru CA^E>T^ tumours more frequently displayed high tumour grade morphology (59.1% grade 3), higher HER2^+^ status (immunohistochemistry 3+ or FISH-confirmed) (31.8%), and higher differentiation (**Supplementary Fig. 15a, b**). Single-cell analysis revealed that in both Stru CA spatial groups, Num CA–positive cells consistently contained more centrosomes in the core than at the edge (**Fig. 6a, b**), indicating a conserved numerical asymmetry. Num CA spatial prevalence was also associated with aggressiveness: Num CA^T>E^ tumours were more often grade 3 (40.6%), had greater nodal involvement (31.2%), and lower HER2^+^ frequency (12.5%) compared with Num CA^E>T^ tumours (28.6%, 7.1%, and 35.7%, respectively; **Supplementary Fig. 15c, d**). Mixed histology was more common in Num CA^T>E^ tumours (12.5%), whereas Num CA^E>T^ tumours were associated with lobular carcinomas (28.6% *versus* 6.3%), a subtype typically linked to slower growth and smaller size. Across both Num CA^T>E^ Num CA^E>T^ groups, centrosomes were significantly larger at the edge than in the core. Notably, Num CA–positive cells with larger centrosomes harboured fewer of them (**Fig. 6c, d**), revealing a robust inverse relationship between centrosome size and number across all spatial groups (**Fig. 6e**). These findings highlight distinct spatial CA patterns associated with tumour subtype and behaviour. Tumours displaying a shift from fewer, larger centrosomes at the periphery to smaller, more numerous centrosomes in the core exhibit features of increased aggressiveness. These spatial patterns, resolved through CenSegNet, provide a proxy for tumour heterogeneity and may reflect the evolutionary trajectory of CA during tumour progression.

**Fig. 6.**
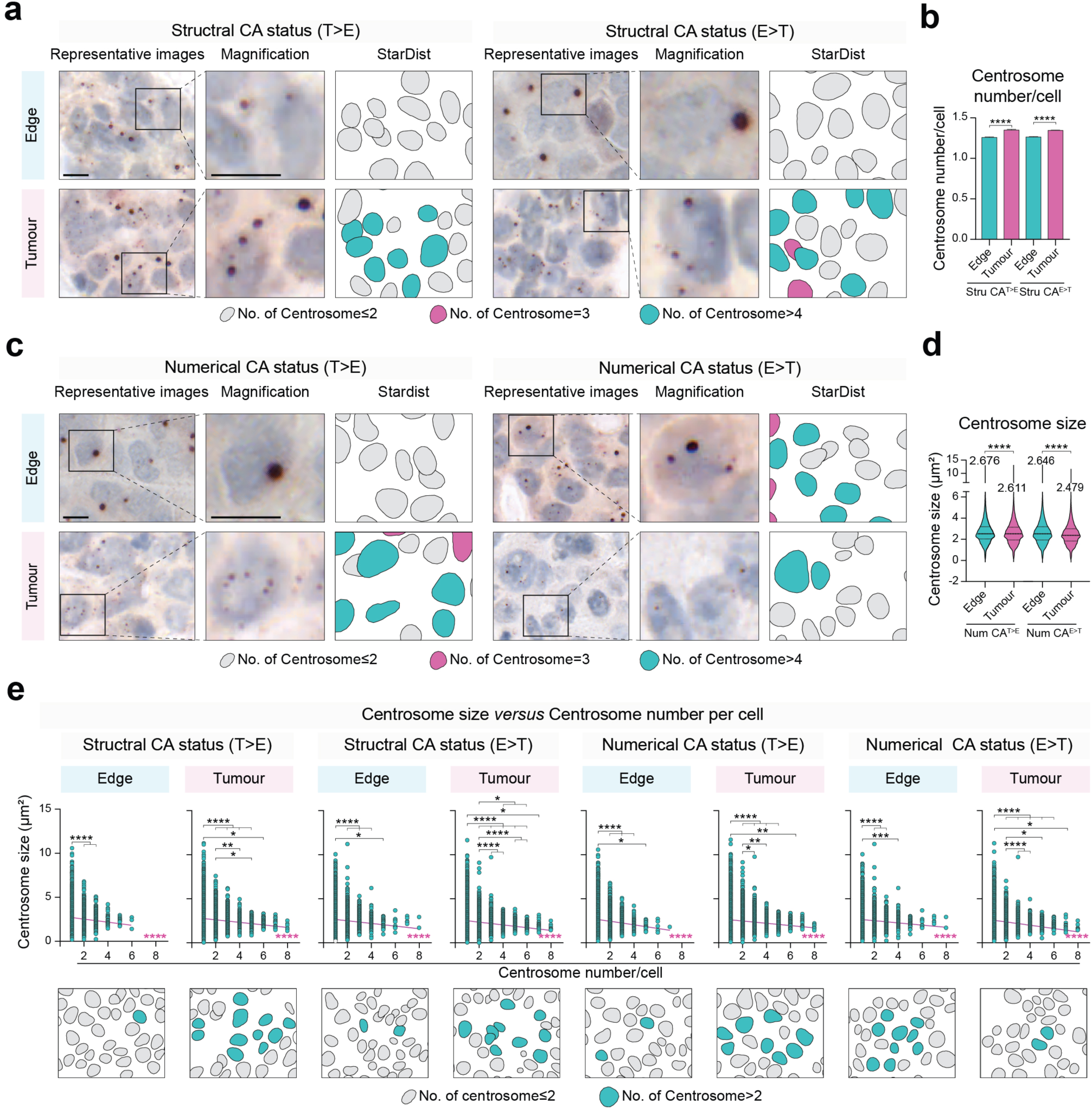
Interplay between Stru CA and Num CA drives centrosome defect spatial heterogeneity in breast edge and tumour regions. **a** Representative immunohistochemistry images of human breast tumour tissues (stained for pericentrin, counterstained with haematoxylin) and corresponding StarDist cell masks showing differences in Stru CA between edge and tumour regions. Left: Stru CA higher in Tumour than Edge. Right: Stru CA lower in Tumour than Edge (teal: cells with ≥4 centrosomes; purple: cells with three centrosomes; grey: cells with ≤2 centrosomes). Scale bars, 10 µm. **b** Comparative analysis of centrosome number per cell in edge and tumour regions for Stru CA^T>E^ and Stru CA^E>T^ groups. Comparisons between Edge and Tumour were performed separately for each condition using two-sided unpaired *t*-test, left: \*\*\*\**P* < 0.0001; right: \*\*\*\**P* < 0.0001. Data are presented as mean ± s.e.m. **c** Representative immunohistochemistry images of human breast tumour tissues (stained for pericentrin, counterstained with haematoxylin) and corresponding StarDist cell masks showing differences in Stru CA and Num CA between edge and tumour regions. Left: Num CA higher in Tumour than Edge. Right: Num CA lower in tumour than edge (teal: cells with ≥4 centrosomes; purple: cells with three centrosomes; grey: cells with ≤2 centrosomes). **d** Comparative analysis of centrosome size in Edge and Tumour regions for Num CA^T>E^ and Num CA^E>T^ groups. Comparisons between Edge and Tumour were performed separately for each condition using two-sided unpaired *t*-test. Left: \*\*\*\**P* < 0.0001; right: \*\*\*\**P* < 0.0001. Data are presented as violin plots showing the distribution of values; dashed lines indicate median and interquartile ranges. **e** Top: correlation between centrosome number and mean centrosome size at the single-cell level in Stru CA^T>E^, Stru CA^E>T^, Num CA^T>E^, and Num CA^E>T^ groups. One-way ANOVA (\*\*\*\**P* < 0.0001 across Stru CA^T>E^, Stru CA^E>T^, Num CA^T>E^, and Num CA^E>T^ groups) with Tukey’s test: Stru CA^T>E^ [Edge: \*\*\*\**P* < 0.0001; Tumour: (top) \**P =* 0.0318, (bottom) \**P* = 0.0416, \*\**P* = 0.0056, \*\*\*\**P* < 0.0001]; Stru CA^E>T^ [Edge: \**P* = 0.0139, \*\*\*\**P* < 0.0001; Tumour: (top) \**P* (2 vs 5) = 0.0362, (top) \**P* (2 vs 6) = 0.0144, \*\*\*\**P* < 0.0001); Num CA^T>E^ [Edge: \**P* = 0.0259, \*\*\*\**P* < 0.0001; Tumour: \**P* = 0.0421, (top) \*\**P* = 0.0081, (bottom) \*\**P* = 0.0025, \*\*\*\**P* < 0.0001]; Num CA^E>T^ [Edge: \*\*\**P* = 0.0003, \*\*\*\**P* < 0.0001; Tumour: (top) \**P =* 0.0340, (bottom) \**P* = 0.0115, \*\*\*\**P* < 0.0001]. Bottom: Corresponding representative StarDist masks showing cells with Stru CA and Num CA in Edge and Tumour regions. Data are presented as individual data points and mean ± s.e.m. Source data are provided as Source Data file.

### CenSegNet integration and accessible deployment for high-throughput centrosome analysis

To support broad adoption and integration into diverse analytical workflows, we provide CenSegNet as both a lightweight application programming interface (API) and an interactive graphical user interface (GUI) (**Fig. 7a, b**). The GUI comprises three modules: a data upload panel for inputting whole-slide or high-resolution images; a prediction module for adjustable inference parameters including probability thresholds and region selection for optimised segmentation of centrosomes and epithelial compartments; and an export tool for structured outputs. Post-inference, users can retrieve per-cell pixel-resolved size estimates, spatial coordinates, and centrosome counts within tissue context (**Fig. 7b)**. The interface supports batch processing and accepts both immunohistochemistry and immunofluorescence formats.

**Fig. 7.**
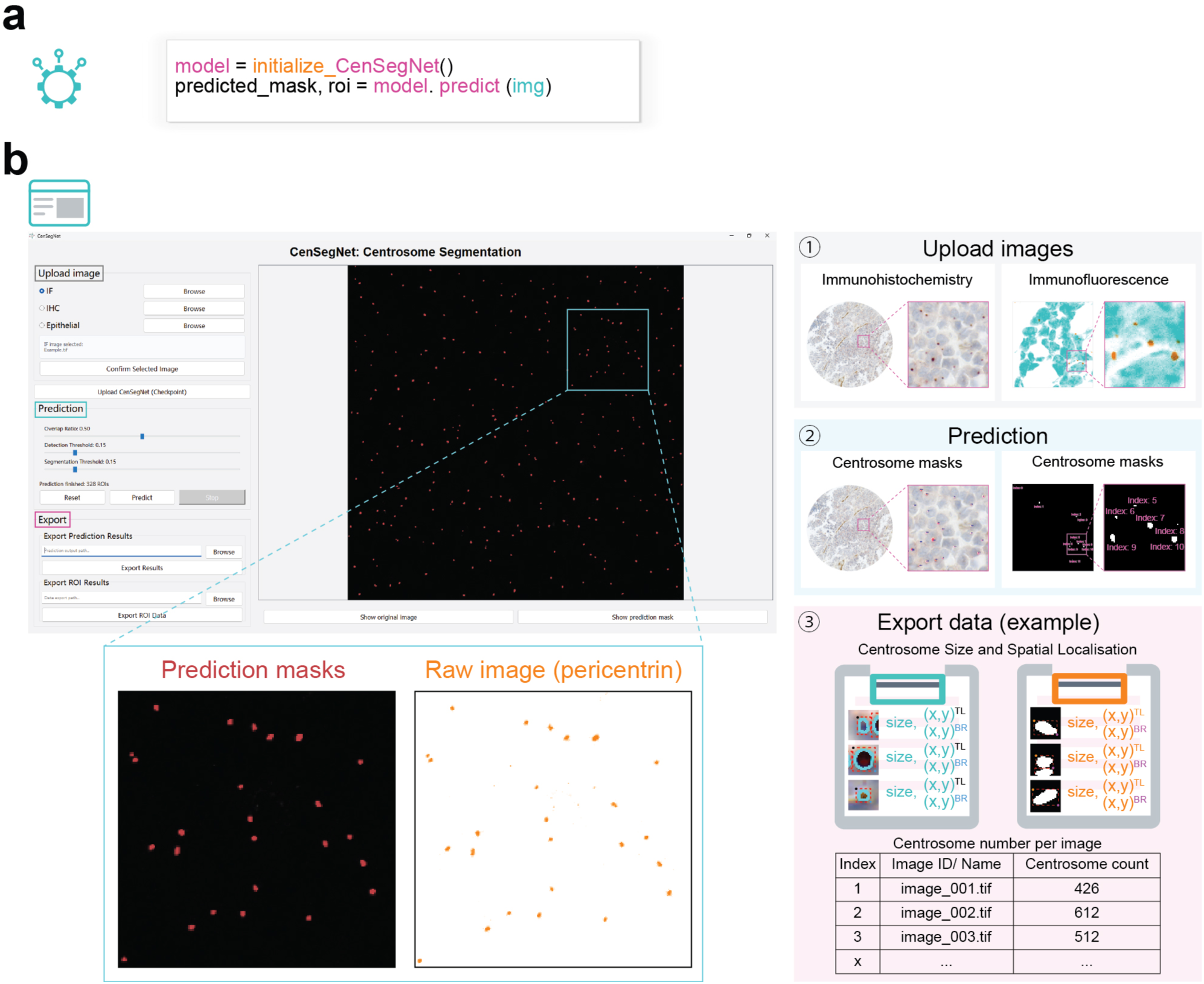
Open accessibility and integration of CenSegNet for broad adoption. **a** CenSegNet can be accessed *via* a Python application programming interface (API). **b** Top: graphical user interface (GUI) of CenSegNet designed for a streamlined workflow, supporting image upload, model-based prediction, and data export. Bottom: interface functions for both immunohistochemistry and immunofluorescence images, enabling users to upload images, apply the relevant prediction models, and export quantitative data including pixel-level centrosome size, localisation, and count.

CenSegNet was executed on a system equipped with four NVIDIA A100 GPUs (40 GB VRAM each), achieving efficient inference across the immunohistochemistry, immunofluorescence, and epithelial datasets. To benchmark prediction time, we evaluated inference on ten randomly selected images from each dataset. Images from the immunohistochemistry and epithelial datasets ranged in dimension from 3,370 × 4,487 to 6,630 × 5,941 pixels, whereas immunofluorescence images were 1,024 × 1,024 pixels. Under these conditions, inference times ranged from 0.28–0.82 min per sample for the immunohistochemistry dataset, 0.19–0.26 min per sample for the immunofluorescence dataset, and 0.24–0.99 min per sample for the epithelial dataset. We next assessed inference performance across three representative hardware configurations, comprising two CPU-only settings and one lower-end GPU configuration (see Methods and **Supplementary Fig. 16**). These comprised a standard office laptop, a higher-performance CPU server, and a freely accessible NVIDIA Tesla T4 GPU available through Google Colab. Specifically, inference performance was evaluated on: (i) CPU #1, a standard consumer-grade office laptop purchased in 2021 and used for routine daily work, equipped with an 11th-generation Intel® Core™ i5-1145G7 processor and 8 GB RAM; (ii) CPU #2, a higher-performance server system equipped with an AMD EPYC™ 9334 32-core processor and 128 GB RAM; and (iii) a GPU configuration using an NVIDIA Tesla T4 with approximately 16 GB VRAM, which is commonly available through Google Colab. CPU #1 was intentionally included to reflect a modest, readily available laptop-class device rather than optimised workstation hardware. Inference on this configuration was fully feasible, requiring approximately 2.6–8.8 min per sample for the immunohistochemistry dataset, 2.1–3.5 min per sample for the immunofluorescence dataset, and 5.8–14.0 min per sample for the epithelial dataset (**Supplementary Fig. 16a**). The corresponding input image dimensions ranged from 3,370 × 4,487 to 6,630 × 5,941 pixels in the TMA cores immunohistochemistry and epithelial datasets, and 1,024 × 1,024 pixels for the immunofluorescence dataset. On the higher-performance CPU server, inference time was substantially reduced to approximately 1.2–3.3 min per sample for the immunohistochemistry dataset, 0.9–1.2 min per sample for the immunofluorescence dataset, and 2.0–5.5 min per sample for the epithelial dataset (**Supplementary Fig. 16b**). Notably, inference on the freely accessible Tesla T4 GPU achieved sub-minute processing per sample across datasets (**Supplementary Fig. 16c**). Although inference speed scaled with available computational resources, these results demonstrate that CenSegNet remains deployable in typical research and clinical-adjacent environments without requiring high-end or specialised hardware.

## Discussion

We introduce CenSegNet, a modular deep learning framework for quantification of centrosomes in epithelial tissues at spatial and single-cell resolution. Unlike previous approaches that treat morphological analysis and segmentation as separate tasks, CenSegNet integrates centrosome detection and phenotyping, nuclear localisation, and epithelial boundary inference into a unified, multichannel pipeline. This enables context-aware segmentation of both structural (Stru CA) and numerical (Num CA) abnormalities across diverse imaging modalities. CenSegNet outperforms established models—including U-Net, SegNet, and DeepLabv3+—in accuracy, generalisability, and morphological fidelity, particularly in densely packed or morphologically heterogeneous tissue regions. The framework comprises three specialised modules: a YOLOv11-based detector trained on over 15,000 annotated centrosomes for robust localisation; a U-Net model for precise segmentation of centrosome area and morphology; and a StarDist-based cell segmentation module optimised for delineating epithelial boundaries in complex tissue architectures. This modular integration, coupled with uncertainty-aware postprocessing, enables systematic and standardised phenotyping of CA subtypes at single-cell and spatial resolution. Applied to 911 sample cores from 127 patients, CenSegNet-based profiling of over 330,000 centrosomes reveals previously uncharacterised spatial trajectories and clinical correlates of Stru CA and Num CA, uncovering their mechanistic uncoupling, age-dependent modulation, and associations with tumour progression, hormone receptor status, HER2 expression, and genomic alterations. While the present cohort is not powered to establish definitive prognostic associations across all CA measures, our analyses identify Stru CA within the tumour core as significantly associated with overall survival and further indicate that spatially resolved numerical and Stru CA patterns align with clinically relevant outcome trends. Together, these findings provide a first indication that specific CA subtypes—particularly Stru CA in defined spatial contexts—may have prognostic relevance and establish a foundation for future studies to rigorously evaluate the independent prognostic value of numerical and structural CA in larger cohorts with longer and more complete clinical follow-up.

Computational tools have been developed previously for CA quantification in epithelial cells. A semi-automated machine learning pipeline quantified PCM defects and numerical CA in immunofluorescence images of normal and breast cancer cells^41^, but the method offers limited spatial resolution and required extensive manual curation. Another semi-automated approach linked centrosome size and number to chromosomal instability in human breast cancer tissues^17^, yet lacked single-cell resolution and scalability. Another pipeline assessed centriole number and length, revealing structural defects arising from fragmentation and ectopic procentriole formation^37^, but without spatially resolved quantification. An automated detection algorithm for high-throughput mapping of CA in ovarian cancer tissues, identified heterogeneous CA phenotypes associated with chromosomal instability and chemotherapy resistance^44^. However, this method did not distinguish between structural and numerical CA or provide single-cell semantic segmentation. CenSegNet addresses these limitations by integrating cellular, spatial, and clinical dimensions of CA across large-scale tissue cohorts. In doing so, it offers new insights into centrosome biology and the functional relevance of CA heterogeneity in breast cancer, with implications for risk stratification and precision oncology. Our orthogonal validation strategy, spanning both immunohistochemistry and immunofluorescence across multiple epithelial tissues and cell culture systems, combined with rigorous manual annotation, modality-specific calibration and sensitivity analyses, enabled CenSegNet to perform consistently across imaging contexts of varying resolution. This approach achieved robust and concordant segmentation of GT335-positive centrioles and pericentrin-labelled pericentriolar material (PCM), reinforcing the intended versatility and generalisability of the framework. Although GT335 chromogenic labelling did not yield reliable or quantifiable signal in paraffin-embedded breast cancer tissues, our validation pipeline demonstrates that CenSegNet remains biologically grounded and fully compatible with pericentrin-based PCM labelling for automated, high-throughput classification of numerical and structural CA in tissue sections. At the same time, we acknowledge that PCM-based labelling alone does not allow direct interrogation of centriole ultrastructural features. This limitation that motivates future complementary studies using electron microscopy, super-resolution microscopy, or expansion microscopy to further resolve centriole-level architectural defects. Consistent with this view, a small number of studies have visualised centriolar markers such as CEP135 and CEP170 in human paraffin-embedded tissues by immunofluorescence, while also recognising the inherently low-throughput nature of these approaches as a key limitation^62, 63^. Beyond breast cancer, CenSegNet has demonstrated generalisability across diverse epithelial tissues—including kidney, colon, and appendix—characterised by high stromal content and architectural complexity (**Supplementary Fig. 17**). To facilitate broad adoption, we provide CenSegNet as an open-source GUI enabling scalable extraction of structured, spatially anchored centrosome metrics. This will allow researchers and clinicians, regardless of computational expertise, to integrate centrosome profiling into histopathological and biomarker discovery workflows. Thus, CenSegNet extends beyond methodological innovation to practical application, accelerating systematic investigation of centrosome biology across anatomically and histologically diverse tissues, and enabling the identification of CA-driven vulnerabilities with potential therapeutic relevance.

Recent studies using a composite centrosome amplification score (CAS) that integrates both numerical and structural abnormalities^7, 43^, showed a progressive increase in CAS from normal breast tissue to invasive carcinoma. However, it did not distinguish the individual contributions of numerical and structural CA. Our spatially resolved single-cell analysis demonstrates that Num CA and Stru CA represent distinct phenotypic axes of centrosome dysregulation. Although they frequently co-occur in tumour tissues, they are uncorrelated at both tissue and single-cell levels and exhibit unique spatial distributions: Stru CA are enriched at tumour edges, while Num CA predominate in tumour cores, suggesting that different regional pressures drive centrosome overduplication *versus* structural enlargement. Single-cell data further uncover a robust inverse relationship between centrosome number and size—cells with multiple centrosomes tend to have smaller ones—indicating a compensatory constraint on total centrosome volume. Consistent with this, centriole over-elongation can induce CA *via* fragmentation and ectopic procentriole formation in breast cancer cells^37^, yet not all centrioles within a cell exhibit these changes, highlighting intra-cellular heterogeneity in elongation susceptibility. Importantly, functional perturbation of PLK4-dependent centriole biogenesis provides experimental support for this proposed mechanistic decoupling. Inhibition of PLK4 activity disrupted the quantitative relationships linking centriole number, centrosome size, and centrosome number, demonstrating that centrosome size can be affected independently of changes in centrosome number. These observations indicate that centrosome structural properties are not solely dictated by numerical amplification and support the existence of separable regulatory mechanisms governing numerical and structural CA under perturbed conditions. Crucially, our data argue against centrosome clustering as the predominant explanation for the enlarged centrosomal structures observed in tumour tissues. Morphometric analyses demonstrate that enlarged centrosomes largely retain high circularity and solidity, consistent with compact, individual centrosomes rather than the irregular, heterogeneous assemblies characteristic of clustered organelles. While centrosome clustering may contribute to a minority of cases, these observations indicate that structural enlargement reflects bona fide centrosome architectural changes rather than unresolved aggregation of multiple centrosomes. Collectively, these findings support a model in which Num CA and Stru CA are mechanistically uncoupled, evolving along orthogonal spatial gradients during tumour progression. This challenges the monolithic view of CA and instead portrays it as a dynamic, regionally modulated process shaped by local microenvironmental cues. The mechanistic decoupling of Num CA and Stru CA has important implications for understanding functional heterogeneity of centrosomal defects in cancer and underscores the need for spatially informed biomarkers.

CenSegNet-based profiling reveals that aging exerts distinct and spatially patterned effects on Stru CA and Num CA in breast cancer. Num CA peaks between ages 50 and 70—overlapping with the menopausal transition and the most common window for breast cancer diagnosis^64–66^—whereas Stru CA accumulates progressively after age 70, indicating a later-life trajectory of centrosome architectural dysregulation. While CA overall increases with age—including in normal tissues—our data suggest that age is not a deterministic initiator but rather a factor that modulates the magnitude and spatial distribution of CA subtypes. In mid-life patients, Num CA is preferentially enriched in tumour cores—regions typically characterised by high proliferation—whereas Stru CA in older individuals extends more diffusely, often into tumour margins, likely reflecting age-associated changes in epithelial architecture, tissue repair dynamics, and microenvironmental stress. A study in prostate cancer has reported elevated CA in patients over 53 years of age^7^. CA increases with age, in normal breast epithelial cells derived from individuals aged 20-80, treated with DNA damage-inducing stimuli^54^. Chronic centrosome overduplication in aging mouse models of intestinal cancer drives aneuploidy and spontaneous tumorigenesis, supporting a role for age-associated CA in early malignant transformation^18^. Centrosome function deteriorates with age, evidenced by accumulation of structural defects and impaired mitotic fidelity^67, 68^. Aging epithelial cells exhibit centrosome fragmentation, aberrant centriole elongation, and altered PCM composition^68, 69^, while broader declines in DNA repair and chromosomal segregation fidelity likely contribute to a permissive environment for CA^68^. Thus, aging tissues accumulating centrosome abnormalities, mitotic fidelity defects, and weakened genomic surveillance, may be more vulnerable to CA-driven tumorigenesis. Our study delineates the divergence of CA subtypes across the lifespan. The inverse scaling between centrosome number and size supports the idea that Num CA and Stru CA act as compensatory, rather than co-occurring, phenotypes. Notably, Stru CA in elderly patients is frequently associated with hormone receptor–negative tumours, implicating centrosome structural dysregulation in the biology of more aggressive or dedifferentiated late-life cancers. CA can induce breast cancer cell dedifferentiation and intrinsically drive high-grade tumours^70^. Age remains a key prognostic factor in breast cancer, influencing tumour subtype distribution, hormone receptor status, and genomic instability^71, 72^. Together, these results establish the first clinically and spatially resolved framework for understanding how aging modulates centrosome biology in human cancer. This framework lays the foundation for developing age-stratified biomarkers and clarifies why specific CA subtypes and their associated chromosomal instability may emerge more frequently or have greater clinical impact at distinct stages of life.

The spatial heterogeneity of CA and its clinical relevance for cancer remain poorly defined^3, 7^. CenSegNet systematically maps Stru CA and Num CA across tumour compartments and stratifies tumours into composite CA subtypes with distinct spatial, biological, and clinical profiles. Stru^high^Num^high^ CA tumours are consistently associated with larger size, greater nodal involvement, and germline genomic alterations. Elevated CA in both core and edge regions suggests a cellular composition primed for proliferative expansion and invasive dissemination. Stru^low^Num^low^ CA tumours often required nodal clearance, supporting evidence that even modest CA can drive aggressive behaviour in permissive genomic contexts—particularly when p53 surveillance is compromised^7, 18, 73^. CenSegNet also uncovers spatial discordance in CA subtypes, with Stru^high^Num^low^ and Stru^low^Num^high^ CA profiles enriched at the invasive front. These spatial signatures are associated with enhanced metastatic potential, consistent with models in which centrosome abnormalities promote invasion through both cell-autonomous^74–76^ and non-cell-autonomous^77–79^ mechanisms. Structural centrosome defects have been shown to drive cell extrusion, facilitating invasion^76, 80^. Our findings corroborate longitudinal studies of tumour progression, such as in Barrett’s oesophagus, where CA appears early in premalignant lesions and expands with p53 loss^73^, supporting a role for CA in tumour initiation rather than as a mere by-product of transformation. Similarly, CA was shown to increase from normal tissue to ductal carcinoma *in situ* (DCIS) to invasive carcinoma, and to correlate with recurrence and poor prognosis^7, 70^. Finally, our observation that Stru CA and Num CA spatial patterns are uncoupled from systemic physiological metrics reinforces CA as a tumour-intrinsic hallmark. Together, these results advance our understanding of centrosome biology in cancer and highlight the power of CenSegNet-driven integration of subcellular organelle features into spatial pathology, with direct implications for clinical decision-making, prognostic modelling, and therapeutic targeting.

By integrating CenSegNet-based centrosome phenotyping with spatially resolved hormone receptor profiling, we uncover compartment-specific associations between CA subtypes and ER/HER2 expression. Num CA is selectively enriched in ER⁻ tumours within the tumour core, whereas HER2⁺ tumours display lower Num CA in the core despite elevated levels at the tumour edge. This spatial divergence suggests that hormone receptor signalling modulates centrosome number and structure in a regionally distinct manner. These observations corroborate previous studies linking CA to hormone receptor status and tumour aggressiveness^37, 42, 75, 81^ and support the hypothesis that ER loss promotes CA through transcriptional or post-translational dysregulation of centriole biogenesis pathways. Basal-like ER⁻PR⁻HER2⁻ breast cancers—characterised by genomic instability and poor prognosis^82^, frequently exhibit high CA, often with centriole fragmentation and ectopic procentriole formation driven by over-elongation^37^. These defects recruit excess PCM, generating supernumerary or structurally abnormal MTOCs that drive mitotic errors and chromosomal instability^83, 84^. In contrast, HER2⁺ tumours are more frequently enriched in discordant CA subtypes (Stru^low^Num^high^ and Stru^high^Num^low^) at the tumour edge, suggesting that HER2 signalling may differentially regulate centrosome number and structure depending on spatial context. This is consistent with evidence that both PLK4 and AURKA expression—key regulators of centriole biogenesis and maturation, respectively—is differentially influenced by HER2 status^81, 85^. Despite high proliferative capacity, HER2⁺ tumours exhibit lower overall CA, raising the possibility that these tumours suppress CA to preserve mitotic fidelity or evade immune detection. Together, our findings identify spatially distinct and mechanistically diverse relationships between hormone receptor signalling and centrosome biology. ER loss is associated with elevated Num CA and mitotic instability in tumour cores, while HER2 signalling appears to exert compartmentalised control over CA subtype distribution. Spatially resolved centrosome profiling thus provides a framework for identifying hormone-specific vulnerabilities that could inform targeted breast cancer therapies.

CenSegNet enables fine-grained stratification of tumours based on the relative abundance of structural and numerical CA, revealing distinct spatial–biological associations. Tumours with Stru CA enriched at the invasive edge (Stru CA^E>T^) are more proliferative, more frequently HER2^+^, and exhibit higher differentiation, whereas those with Stru CA more abundant in the core (Stru CA^T>E^) are more commonly associated with lobular histology and slower growth. Single-cell analyses further show that tumours with grade 3 histology, nodal involvement, and mixed subtypes—hallmarks of aggressive disease—are more prevalent in the Num CA^T>E^ subgroup, while Num CA^E>T^ tumours are enriched for lobular carcinomas, typically linked to indolent behaviour^82^. These spatially resolved patterns corroborate previous reports suggesting that numerical centrosome abnormalities increase with tumour progression and are more frequent in aggressive basal-like carcinomas^7, 37^. Our single-cell data also reveal an inverse relationship between centrosome number and size—where edge regions harbour fewer but larger centrosomes—supporting a model in which Stru CA at the periphery primes cells to acquire invasive behaviours, while Num CA in the core drives proliferation and genomic instability. This dynamic interplay likely reflects microenvironmental influences on CA trajectories during tumour progression. These findings extend the “CA set point” concept^86^, which postulates that tumours maintain a context-dependent equilibrium of CA phenotypes to balance proliferation, invasion and survival. CenSegNet-based spatial profiling demonstrates that CA is not only subtype-specific but also spatially regulated, providing new insights into the architectural evolution and heterogeneity of breast cancer.

Although evaluation of the relevance of CA patterns to patient outcome in breast cancer did not identify Num CA and Stru CA as statistically significant independent predictors of overall or recurrence-free survival following multivariable adjustment in the present cohort, several consistent and biologically meaningful patterns were observed. Notably, lower levels of structural CA within tumour cores were significantly associated with more favourable overall survival, whereas tumours characterised by higher structural CA coupled with lower numerical CA in the core tended to exhibit less favourable outcomes. These patterns closely mirror the clinicopathological associations identified throughout this study, including tumour size, nodal involvement, and family history, supporting the view that CA subtypes reflect integrated tumour states rather than acting as isolated prognostic variables. These observations should be interpreted in the context of cohort size and follow-up duration. The BeGIN cohort is a relatively recent study, and the limited number of outcome events constrained statistical power and model stability, particularly in multivariable analyses. Within these constraints, additional exploratory stratification analyses revealed internally coherent outcome trends, providing further support for the clinical relevance of the orthogonal and spatial organisation of numerical and structural CA. Consistent with our broader findings, these results suggest that elevated structural CA—particularly when enriched within tumour cores—may mark more aggressive tumour behaviour, whereas numerical CA in relative isolation appears better tolerated. Validation in larger, cancer-specific cohorts with longer and more complete clinical follow-up will be required to determine whether these CA subtypes provide independent prognostic or predictive value beyond established clinicopathological determinants. Importantly, our breast cancer-specific observations are consistent with a growing body of evidence across epithelial malignancies implicating CA as clinically informative features of tumour progression. Pan-cancer analyses have linked centrosome amplification–associated transcriptional signatures to genomic instability and aggressive tumour phenotypes across diverse cancer types^87^, while tumour-specific studies in colorectal, prostate, bladder, and ovarian cancers have demonstrated robust associations between centrosome defects, chromosomal instability, tumour grade, and disease progression^10, 88–90^. Together, these findings position numerical and structural CA as biologically grounded and clinically relevant features that integrate with tumour context rather than serving as standalone prognostic markers. By enabling scalable, spatially resolved quantification of CA subtypes, CenSegNet provides a framework for dissecting how centrosome dysregulation intersects with tumour architecture and clinical behaviour, and lays the foundation for future studies evaluating CA-associated vulnerabilities in precision oncology.

In summary, CenSegNet delivers the first fully integrated and spatially resolved framework for profiling CA at single-cell resolution across large-scale human cancer tissues. By enabling precise, high-throughput quantification of both numerical and structural CA phenotypes, CenSegNet uncovers distinct mechanistic, temporal, and spatial trajectories of centrosome dysregulation in breast cancer. The discovery that numerical and structural CA are decoupled—not only in cellular architecture but also in their associations with clinical features such as tumour grade, hormone receptor status, germline mutation, and patient age—advances our understanding of how centrosome abnormalities contribute to intratumoral heterogeneity and tumour progression, with considerable implications for therapeutic resistance. Importantly, the identification of compensatory dynamics between centrosome number and size, and their divergent distributions across tumour cores and margins, points to context-specific roles in modulating local tumour ecology and genomic instability. These insights challenge the longstanding view of CA as a uniform driver of malignancy, instead positioning CA subtypes as distinct functional modules in tumour evolution. CenSegNet thus provides a foundation for developing CA-based biomarkers to stratify patients by tumour subtype, age, and aggressiveness, and opens opportunities for therapies targeting CA-driven vulnerabilities. Given the availability of PLK4, AURKA, and HSET inhibitors^91–95^, spatial CA maps could guide personalised strategies, particularly in tumours with discordant CA phenotypes. Future studies integrating CenSegNet-based CA profiling with transcriptomic and proteomic analyses will be essential to uncover the molecular drivers of spatial CA dynamics and clarify their roles in tumour progression, metastatic dissemination, and therapy resistance.

## Methods

### Ethics and human breast tissues

The study participants were a subgroup of women diagnosed with early breast cancer who were recruited to a single-centre prospective observational cohort study at University Hospitals Southampton, “Investigating Outcomes from Breast Cancer: Correlating Genetic, Immunological and Nutritional Predictors (<u>BeGIN</u>)^96, 97^.” All procedures performed in studies involving human participants were in accordance with the ethical standards of the institutional and/or national research committee and with the 1964 Helsinki declaration and its later amendments or comparable ethical standards. All participants in BeGIN gave written informed consent. The research ethics committee approved the study (Research Ethics Committee (REC) - Cambridgeshire and Hertfordshire reference number: 14/EE/1297). Women were eligible for the BeGIN study if they were aged >18 years and diagnosed with invasive breast cancer or DCIS at University Hospital Southampton after May 2015. Linked anonymised patient information, including patient characteristics, tumour characteristics and clinical management, were extracted from the hospital electronic record system. Body composition parameters were measured using Bioelectrical Impedance Analysis (BIA) with a phase-sensitive, 8-electrode device (Seca mBCA515)^98^. To conduct this study, 911 cores from 127 breast cancer patients were used. The TMAs were constructed from formalin fixed paraffin embedded (FFPE) histopathology tissue blocks from surgical treatment surplus to diagnostic requirements. Colon, kidney, and appendix tissue were incorporated into breast tissue TMAs to facilitate orientation during sectioning and analysis. Colon and kidney samples consisted of histologically normal tissue, the status of which was independently verified by a board-certified pathologist. A summary of the clinicopathological information linked to the human breast samples used in this study is included in **Supplementary Table 1**. Anonymous data from the BeGIN study is available for request to researchers who provide a completed Data Sharing request form that describes a methodologically sound proposal, for the purpose of the approved proposal. Proposals will be reviewed by the study steering committee. Data will be shared once all parties have signed relevant data sharing documentation, covering the study steering committee conditions for sharing and if required, an additional Data Sharing Agreement from the Sponsor.

### Ethics and mice

BALB/c HER-2/neu transgenic mice (referred to as BALB-NeuT)^99^ carrying the transforming rat *Her-2/neu* oncogene under control of a MMTV-LTR were used. All experimental procedures involving mice were approved by the University of Southampton Local Ethics Committee and registered with the Ethics and Research Governance Online II (ERGO II; ID: 65385). All animal work was conducted in accordance with UK Home Office regulations, adhering to the principles of the 3Rs (Replacement, Reduction, Refinement) and the ARRIVE (Animal Research: Reporting of In Vivo Experiments) guidelines to minimise animal suffering throughout the study. Mice were housed in a specific pathogen-free (SPF) facility under controlled environmental conditions, including regulated temperature and humidity, with a 12-hour light/dark cycle. Animals had ad libitum access to standard chow and water.

### Cell culture

MCF10A is a non-transformed human mammary epithelial cell line (ATCC® CRL-10317). The MCF10A-PLK4 cell line is a genetically engineered derivative of MCF10A that enables inducible overexpression of Polo-like kinase 4 (PLK4), a master regulator of centrosome duplication whose upregulation induces centrosome amplification^75^. The MCF10A-PLK4 and MCF10DCIS cell lines were kindly provided by Professor Susana Godinho (The Barts Cancer Institute, Queen Mary University of London). All cells were cultured in DMEM/F12 medium (Invitrogen), supplemented with 10% donor horse serum (Gibco, 31331028), 20 ng/ml human epidermal growth factor (EGF; Sigma, E9644), 10 μg/ml insulin (Sigma, I1882), 100 μg/ml hydrocortisone (Sigma, H0888), 1 ng/ml cholera toxin (Sigma, C8052), and 50 U/ml penicillin with 50 μg/ml streptomycin (Life Technologies). Cells were maintained at 37 °C in a humidified incubator with 5% CO₂. To induce PLK4 overexpression, cells were treated with doxycycline (Sigma, D9891) at 2 μg/ml for 48 hours. To inhibit PLK4, cells were treated with 500 μM Centrinone (Tocris Bioscience, 5687) for 5 days.

### Tissue microarray construction and immunohistochemistry

Tissue microarrays (TMAs) were constructed using formalin-fixed, paraffin-embedded (FFPE) tissue samples obtained following breast cancer surgery from 127 patients diagnosed with primary invasive breast carcinoma at University Hospital Southampton between July 9, 2015, and January 31, 2019, and participating in the BeGIN study. All patients underwent standardised treatment at a single institution, consisting of surgery, followed by adjuvant treatments according to local and national protocols. Pathological evaluation of hormone receptor and HER2 expression was conducted according to established national and international guidelines^100, 101^. Immunohistochemistry was used to determine hormone receptor status, and in situ hybridization (ISH) was employed to confirm HER2 positivity for tumours with an IHC score of 2+. All procedures were performed within the standard clinical diagnostic pathway. A total of 911 sample cores were systematically sampled from three distinct, pathologically classified regions for each patient: tumour tissue (Tumour), tumour margin (Edge), and tumour-free tissue (Normal). Each patient represented by three technical replicate cores from Tumour, Edge, and Normal regions, were procured for analysis. Each 0.6 mm in diameter and 5 µm in thickness were extracted from formalin-fixed, paraffin-embedded specimens and arrayed into recipient tissue microarray (TMA) blocks. This tri-regional, triplicate-core sampling strategy was designed to provide a comprehensive and robust representation of the tissue heterogeneity within and around the tumour microenvironment. TMA sections were mounted onto TOMO® adhesion microscope slides. Immunohistochemistry was performed using the Dako Autostainer Link 48 automated platform. Endogenous peroxidase activity was quenched using EnVision FLEX blocking reagent (Dako), followed by a 30-minute incubation with the primary antibody against pericentrin (1:500 dilution; Abcam, ab4448). Signal amplification and enzymatic detection were achieved using EnVision FLEX HRP (Dako, 20 minutes) and Rabbit Link (Dako, 15 minutes). Slides were counterstained with haematoxylin following three 5-minute washes in 3-amino-9-ethylcarbazole (AEC).

### Assessment of hormone receptor status

ER or PR expression were evaluated by immunohistochemistry and scored using the Allred system (range 0–8). Tumours with an Allred score ≥ 3 were classified as ER or PR positive. HER2 status was determined by immunohistochemistry and scored as 0, 1+, 2+, or 3+. Tumours with a score of 3+ were classified as HER2 positive. Cases with a score of 2+ were considered equivocal were further assessed by fluorescence in situ hybridization (FISH). Tumours with FISH amplification were designed as HER2 positive, defined as a HER2/CEP17 ratio ≥ 2.0 or average HER2 copy number ≥ 6.0 signals per cell.

### Immunofluorescence

The following primary antibodies were used: anti-GT335 (1:800; Adipogen, AG-20B-0020-C100), anti-pericentrin (1:250; Abcam, ab4448), and anti-keratin 8/18 (KRT8/18; 1:300; Origene, BP5007). Secondary antibodies (Life Technologies) included goat anti-mouse (A-32723), anti-rabbit (A-11037 and A-11008), and anti-guinea pig (A-21450), each conjugated to Alexa Fluor 488, Alexa Fluor 594, or Alexa Fluor 647, and used at a final concentration of 5 μg/ml.

OCT-embedded mammary gland sections (30 µm thick) from BALB-NeuT mice were cryosectioned, air-dried for 30 minutes, and fixed in 4% paraformaldehyde (PFA) for 20 minutes at room temperature. Sections were permeabilised for 45 minutes with 0.1% Triton X-100 in PBS, then blocked for 2 hours in a solution containing 2% bovine serum albumin (BSA), 5% foetal bovine serum (FBS; Gibco), and 0.1% Triton X-100 in PBS. Sections were incubated overnight at 4 °C with primary antibodies against pericentrin, GT335, and KRT8, followed by washing and incubation with the appropriate secondary antibodies for 2 hours at room temperature. Nuclei were counterstained with DAPI using Fluoroshield mounting medium (Sigma, F6057).

MCF10A and MCF10A-PLK4 cells were fixed in anhydrous methanol at −20 °C for 10 minutes, followed by permeabilisation with 0.1% Triton X-100 in PBS for 2 minutes. Cells were then washed three times for 5 minutes each with 0.1% Triton X-100 in PBS. Blocking was performed using 3% BSA in 0.1% Triton X-100 in PBS for 1 hour at room temperature. Cells were incubated overnight at 4 °C with primary antibodies against pericentrin and GT335. After washing, cells were incubated with the appropriate secondary antibodies for 1 hour at room temperature and counterstained with DAPI using Fluoroshield.

### Microscopy and image annotation

Immunohistochemistry images of TMA cores were acquired using a Zeiss Axio Imager Z1 upright microscope (Zeiss) equipped with an AxioCam MRc5 colour camera. Whole-slide images were captured using Zeiss ZEN imaging software according to a predefined whole-slide brightfield scanning protocol (Whole Slide [WS] Brightfield [BF] fold-light) and using a 20× objective lens. The scaling per pixel was 0.22 × 0.22 μm. This imaging configuration enabled high-resolution acquisition of tissue sections with consistent illumination and contrast across samples. Individual TMA core images, ranging from approximately 3000 × 3000 to 8000 × 8000 pixels, were subsequently exported from the scanned whole slides for downstream analysis. Differences in image dimensions reflect variation in exported tissue area rather than differences in imaging resolution.

Immunofluorescence images were captured using an inverted STELLARIS 5 Confocal Laser Scanning Microscope (Leica Microsystems), equipped with a 40× Glycerol immersion objective: HC PL APO 40×/1.25 GLYC CORR CS2, (Leica Microsystems, Germany). 16-bit micrograph images and Z-stacks were acquired using voxel sampling in excess of Nyquist requirements (Rayleigh resolution/6 and 0.2 μm Z-step size respectively), using fields of view (FOVs) ranging from 1024 × 1024 to 2048 × 2048 pixels. The confocal pinhole was set to 0.5 Airy units, and images were reconstructed using the Leica Lightning computational super-resolution algorithm (Leica Microsystems), yielding an effective lateral resolution of approximately 120 nm. All cells from all FOVs obtained during the experiments were included in the evaluation of model performance. Image processing was performed using Fiji software (https://imagej.net/software/fiji/)^102^.

Manual annotations of centrosomes and epithelial regions were performed by Jiaoqi Cheng (see **Supplementary Fig. 18**), under the supervision of two domain experts, Dr Constantinos Savva (SCE, Medical Oncology) and Dr Margaret Ashton-Key, a nationally and internationally recognised Haematopathologist, both with extensive clinical experience, and were subsequently reviewed by these experts. Centrosome size quantification in immunohistochemistry images was performed using calibrated spatial resolution, wherein each pixel corresponded to an area of 0.0483 μm^2^. For each centrosome, pixel-level segmentation masks were used to compute total pixel occupancy, which was then multiplied by the calibrated pixel area to derive centrosome size in μm^2^. To further ensure annotation robustness, a subset of annotations was independently evaluated by additional observers with complementary expertise in cell biology (Mengyang Gou) and computational analysis (Peixin Zuo). Inter-observer agreement was quantified using intersection-over-union (IoU), F1 score, and Cohen’s κ coefficient. Discrepancies between annotators were systematically assessed and resolved through consensus discussion with the senior experts, ensuring consistency and high-quality annotation standards across the dataset.

Cohen’s k was calculated as:

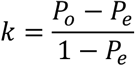

With:

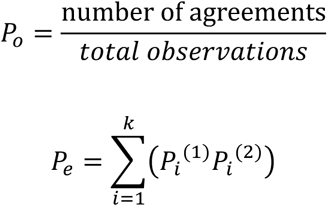

where *P_o_* represents observed agreement between annotators, *P_e_* represents expected agreement by chance.

To generate the immunohistochemistry training dataset, images from 78 patients were included, comprising human breast cancer tissue, normal breast tissue, normal liver, and normal kidney samples. Within these images, annotations were made for centrosomes (n = 14,679), epithelial compartments (n = 2,486), and stromal compartments (n = 108), using QuPath v0.5.1 (https://qupath.github.io/)^103^. All centrosome annotations were performed under 200× magnification for each region of interest (ROI). The manual annotation process required over 200 person-hours. Annotations were exported as GeoJSON files.

For the immunofluorescence training dataset, only 15 high-resolution image z-stacks of MCF10A and MCF10A-PLK4 cell lines were required. Annotation was performed using Cellpose 3.0^104, 105^. Datasets were first maximum-intensity projected, then split into contrast-adjusted single-channel images, with boundaries defined by edge features visible in the blue channel. The manual annotation process took approximately 16 person-hours. All annotations were exported as SVG files.

### Measurements of centrosome morphometric descriptors

Morphometric features of annotated centrosomes were quantified using QuPath (version 0.5.1). For each centrosome, geometric descriptors were computed from the region of interest (ROI) boundary.

Circularity was calculated as:

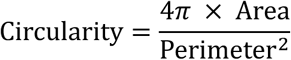

Circularity is a metric describe how close shape is to a circle, providing a normalized measure of shape compactness with a value of 1 corresponding to a perfect circle and decreasing with increasing boundary irregularity.

Solidity was calculated as:

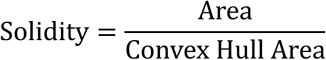

Solidity is a metric describe how convex (non-indented) the shape is reflecting the degree of concavity within the ROI, where values approaching 1 indicate fully convex shapes. Solidity was calculated as the ratio of the annotation area to the area of its convex hull. These metrics were derived directly from ROI geometries within QuPath using its built-in measurement functions without additional post-processing.

### Image processing and model training

**Image processing.** For immunohistochemistry image processing in the CenSegNet pipeline, raw microscopy images were pre-processed to enhance contrast and suppress background noise, thereby improving the visibility of cellular structures. Each image was cropped into overlapping patches of 256 × 256 pixels with a stride of 300 pixels. Patches exhibiting artefacts or poor quality were manually excluded to ensure dataset integrity. The final immunohistochemistry dataset comprised 1,122 annotated patches, each containing labelled information on centrosome location and size. For immunofluorescence images, RGB channels were converted to greyscale to reduce dimensionality and simplify the training process. This transformation allowed the model to remain invariant to colour information while improving computational efficiency.

For immunohistochemistry image processing in the CenSegNet pipeline, raw microscopy images were pre-processed to enhance contrast and suppress background noise, thereby improving the visibility of cellular structures. Each image was cropped into overlapping patches of 256 × 256 pixels using a stride of 300 pixels. Patches containing technical artefacts, low signal quality, or tissue damage were manually excluded to ensure dataset integrity. The final immunohistochemistry training dataset comprised 1,122 annotated patches, each containing labelled information on centrosome location and size. To improve robustness to staining variability and tissue heterogeneity, data augmentation strategies—including hue, saturation, and value (HSV) colour jitter and geometric transformations—were applied during training. Immunofluorescence images exhibit signal characteristics that differ fundamentally from immunohistochemistry, including fluorescence-dependent emission, broader dynamic range, and channel-specific intensity variability. To account for these properties while preserving biologically relevant structural information, a modality-specific preprocessing pipeline was implemented. Immunofluorescence images were similarly cropped into overlapping 256 × 256-pixel patches, and RGB channels were converted to greyscale prior to training. This conversion reduced inter-channel redundancy and sensitivity to channel-specific intensity fluctuations while retaining the spatial features required for centrosome detection, thereby improving computational efficiency and model invariance to colour information.

Together, these modality-specific preprocessing and calibration steps ensured that CenSegNet received appropriately normalised inputs tailored to each imaging modality. This strategy minimised modality-driven bias, improved generalisability across immunohistochemistry and immunofluorescence datasets, and enabled consistent centrosome detection and quantification across imaging contexts with distinct signal generation and noise characteristics.

**Model architecture and training.** CenSegNet employs a modular three-step architecture comprising approximately 40 million trainable parameters. The first step consists of a detection head that localises candidate ROIs, while the second stage performs fine-grained segmentation within these ROIs to achieve precise spatial delineation. This decoupled design enables task-specific optimisation and improves memory efficiency and training stability. In the detection stage, we fine-tuned YOLOv11-seg model^45^ (https://github.com/ultralytics/ultralytics) using our own training set, guided by a composite loss function comprising box loss, segmentation loss, classification loss, and distribution focal loss. These components respectively optimise object localisation, foreground-background separation, class prediction, and robustness to complex spatial distributions. Training was performed using the AdamW optimiser (learning rate = 0.002, momentum = 0.9) with a batch size of 16 over 300 epochs. The segmentation step 2 employed a U-Net architecture^46^ with three input channels (RGB) and one output channel. Given the small size of centrosomes, segmentation was formulated as a binary classification task. The model was trained for 100 epochs using the Binary Cross Entropy with Logits loss function (BCEWithLogitsLoss). Optimisation was performed using RMSprop (learning rate = 1 × 10^-4^, weight decay = 1 × 10^-8^, momentum = 0.9), which stabilised training and improved convergence for small target structures. Training was conducted in Python (v3.10) using PyTorch (v2.1). We applied data augmentations, such as HSV colour jittering and geometric transformations, which can be easily implemented using the torchvision.transforms module in PyTorch. The detection head was trained independently for approximately 24 hours, followed by an additional 12 hours of training for the segmentation head. All experiments were run on four NVIDIA A100 GPUs (40 GB VRAM each) using PyTorch’s Distributed Data Parallel (DDP) framework with NVIDIA Collective Communications Library (NCCL) backend for gradient synchronisation. Input batches were evenly partitioned across GPUs, with local gradient computation and synchronisation via NCCL’s optimised collective communication, achieving near-linear scaling in throughput. Batch sizes were dynamically adjusted to maximise GPU utilisation while maintaining training stability.

**Sensitivity analysis of detection and segmentation thresholds.** We performed sensitivity analyses to assess the effect of detection and segmentation threshold selection across representative imaging conditions. Four image types were included: immunohistochemistry, confocal immunofluorescence, Lightning super-resolution immunofluorescence, and haematoxylin-stained images. For each image type, threshold values ranging from 0.1 to 0.8 were evaluated. In one analysis, the detection threshold was varied while the segmentation threshold was fixed at 0.15 (default value). In a second analysis, the segmentation threshold was varied while the detection threshold was fixed at 0.15. Model performance was quantified at each threshold using the F1 score, IoU, and Cohen’s κ, and results were compared across threshold values and image types.

**Inference-time performance and computational requirements.** Inference performance was evaluated on three representative hardware platforms: a consumer-grade laptop CPU (Intel Core i5-1145G7, 8 GB RAM), a high-performance server CPU (AMD EPYC 9334, 32 cores, 128 GB RAM), and an NVIDIA Tesla T4 GPU with ∼16 GB VRAM, corresponding to the standard free-tier Google Colab GPU. The laptop configuration was included to assess feasibility on modest, readily available local hardware, whereas the server CPU and GPU represented progressively higher-performance computing environments. Runtime was measured as minutes per sample across the immunohistochemistry, immunofluorescence and epithelial datasets. The corresponding input image resolutions were 3370 × 4487 to 6630 × 5941 pixels for the immunohistochemistry and epithelial datasets, and 1024 × 1024 pixels for the immunofluorescence dataset.

**Comparative segmentation models.** To benchmark CenSegNet, we compared its performance against established segmentation models including U-Net^46^, SegNet^48^, and DeepLabv3+^49^. Each model was trained on uniformly sized 256 × 256 image patches cropped from the original dataset. Official implementations were used without architectural modifications. Comparative experiments were conducted with and without ImageNet pretraining to assess the impact of transfer learning. Model outputs were evaluated against ground truth segmentation masks using a composite loss function combining weighted binary cross-entropy and Dice loss, balancing pixel-wise accuracy with regional overlap. All models were trained under identical conditions: Adam optimiser (initial learning rate = 1 × 10^-4^), cosine learning rate decay, L2 regularisation (weight decay = 1 × 10^-8^), batch size of 16, and early stopping based on validation loss plateauing. A total of 108 immunohistochemistry images were manually annotated and divided into training (n=80), validation (8), and test (n=20) datasets. An additional private dataset comprising 25 images was generated and manually curated for evaluation.

**Cell segmentation.** For cell nucleus segmentation step 3, we integrated StarDist^47^ with the QuPath platform. StarDist offers state-of-the-art performance in dense and noisy biological environments. Following accurate nucleus segmentation, cell membrane boundaries were estimated using the spatial coordinates of nuclei as anchor points. Expansion thresholds of 3, 4, 5, and 6 μm were tested and validated against KRT8-stained images to determine the optimal value for our dataset.

## Evaluation metrics

**Centrosome segmentation performance.** Centrosome segmentation performance was evaluated on a test set comprising 25 immunohistochemistry and 17 immunofluorescence images. Metrics included precision, recall, intersection over union (IoU), F1 score. IoU was defined as the common area between the predicted segmentation and the ground truth:

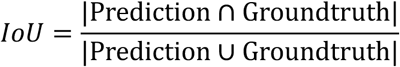

The F1 score, representing the harmonic mean of precision and recall, was calculated as:

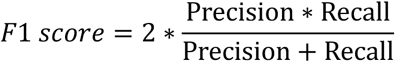

With:

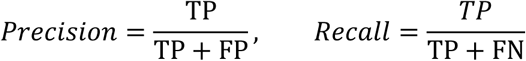

Where, TP, FP, and FN denote true positives, false positives, and false negatives, respectively. Both IoU and F1 scores range from 0 to 1, with higher values indicating superior segmentation performance. These metrics provide a comprehensive assessment of spatial overlap, agreement, and pixel-wise accuracy, with higher values indicating better performance in medical image segmentation.

**Epithelial segmentation performance.** To assess the performance of epithelial segmentation by CenSegNet, we quantified the F1^IoU50^ metric^106^, a widely adopted benchmark in biomedical image analysis. IoU50 means the IoU threshold was set to 0.5 (or 50%). A predicted bounding box was considered a correct detection if: IoU with ground truth ≥0.5.

### Survival analysis

Among 98 patients, 10 deaths occurred during follow-up. Median follow-up was 90.64 months. The 3-year overall survival was 97.96%, and 5-year overall survival was 95.92%. Overall survival (OS) was defined as the interval from the date of diagnostic biopsy to death from any cause or last follow-up. Recurrence-free survival (RFS) was defined as the interval from the date of diagnostic biopsy to the first documented recurrence, death from any cause or last follow-up, whichever occurred first. Survival probabilities were estimated using the Kaplan–Meier method and compared between groups using the log-rank test. Multivariable Cox proportional hazards regression was performed to evaluate whether the proposed CA status was independently associated with OS and RFS. Variables included in the multivariable models were selected based on clinical relevance and/or previously reported prognostic significance. High and low Stru CA and Num CA status were defined separately using the respective median CA values as cutoffs. Hazard ratios (HRs), 95% confidence intervals (CIs) and P values were reported.

### Statistical analysis

The exact n is stated in the corresponding figure legend. GraphPad Prism 10.3.1 (GraphPad Software) was used to perform statistical significance analysis. Normality and lognormality were assessed prior to statistical analysis. For datasets exhibiting a normal distribution, comparisons between two groups were conducted using an unpaired t-test, while comparisons across multiple groups were performed using one-way analysis of variance (ANOVA), followed by Tukey’s post hoc test for multiple comparisons or two-way ANOVA, followed by Tukey’s post hoc test for multiple comparisons. For datasets that deviated from normality, non-parametric testing was employed, using the Kruskal-Wallis test for multiple group comparisons, followed by the Mann-Whitney U test as a post hoc analysis. For correlation analysis, a two-tailed Pearson test was used followed by simple linear regression for graphical representation for correlation analysis. To compare the composition of groups based on categorical variables (e.g., histological tumour grade, T stage, histological tumour types, hormone status, nodal status, nodal clearance, HER2 status), Fisher’s exact test was employed. This non-parametric test was chosen for its appropriateness with count data and its ability to provide accurate p-values when expected cell counts are low (<5). Survival analyses were performed using the Kaplan–Meier method, and differences between groups were assessed using the log-rank test. All comparisons were two-sided. All values were presented as mean ± s.e.m. For all statistical tests, \**P* ≤ 0.05, \*\**P* ≤ 0.01, \*\*\**P* ≤ 0.001 and \*\*\*\**P* ≤ 0.0001 were considered significant.

## Assistance with manuscript preparation

During the preparation of this manuscript, ChatGPT (OpenAI, version 4) was used to assist with stylistic and grammatical refinement. All AI-generated content was critically reviewed and edited by the authors, who take full responsibility for its accuracy and integrity.

## Data availability

Examples of immunohistochemistry and immunofluorescence image datasets used for benchmarking and for testing CenSegNet within the demo version of the pipeline are available on https://github.com/SKELab/CenSegNet/ and https://zenodo.org/records/17131573. All other relevant data supporting the key findings of this study are available within the article and its Supplementary Information files or upon reasonable request. A summary of the human breast samples used in this study are included in Supplementary Table 1. For participants on the BeGIN study, further donor anonymised clinicopathological information is available upon reasonable request, provided all relevant ethics approvals are in place (see “Ethics and human breast tissues” section for further details). The source data that support the findings in all Figures and Supplementary Figures are provided as a Source Data file within the paper. All reagents generated in this study are available upon reasonable request.

## Code availability

The source code and software CenSegNet as a ready-to-use executable with a quickstart guide, example datasets and step-by-step procedures are freely available at https://github.com/SKELab/CenSegNet/ and https://zenodo.org/records/17131573.

## Acknowledgements

We would like to thank Dr. Mark Willett at the Imaging and Microscopy Centre for valuable assistance with fluorescence microscopy; Monette Lopez at the Research Histology Facility, University Hospital Southampton for performing immunohistochemistry staining and imaging; Dr Margaret Ashton-Key for classifying breast tumour compartments (normal, edge, core) and providing advice on immunohistochemistry staining conditions; Professor Susana Godinho for providing the MCF10A and MCF10A-PLK4 cell lines. In addition, we thank Dr. Xi Jia, Dr. Colinda Scheele, Dr. Marcin Przewloka, and Dr. Kif Liakath-Ali, for critically reviewing the manuscript. Cartoons in Figs. 1a, 1c and Supplementary Fig. 5a and 18c were created using https://www.freepik.com/. This work was supported by an MRC New Investigator Research Grant (MR/R026610/1) awarded to S.E. and an Institute for Life Sciences Pilot Grant awarded to R.C., S.B. and S.E. JC was supported by the Gerald Kerkut Charitable Trust and a China Scholarship Council (CSC) PhD studentship. The BeGIN study was supported by Breast Cancer Now. We would like to acknowledge Dr. Keqiang Fan, who sadly passed away during the preparation of this manuscript. Dr. Fan contributed to the early development of CenSegNet, providing an important foundation for the work presented here. We are grateful for his valuable collaboration and the opportunity to have worked with him, and we remember him with deep respect and appreciation.

## Author contributions

J.C. designed and performed experiments, analysed, interpreted the data, and wrote the manuscript. K.F. trained models and developed CenSegNet. C.S. M.B and X.D. supervised centrosome annotation and performed transfer annotations from GeoJSON Files to PNG files and data analysis. E.R. and M.G. performed immunofluorescence experiments. M.G. and P.Z. performed validation of the centrosome and epithelial annotations by comparison with annotations generated by J.C.. R.C. provided ethically approved BeGIN breast cancer tissues. S.B. supervised J.C. and interpreted the data. X.C. supervised K.F. on CenSegNet development. S.E. conceived and designed the project, analysed and interpreted the data, supervised J.C., wrote the manuscript, and acquired funding. All the authors provided intellectual input, edited, and approved the final manuscript.

