## Supplementary Figures for "CenSegNet: a generalist high-throughput deep learning framework for centrosome phenotyping at spatial and single-cell resolution in heterogeneous tissues"

Cheng *et al.*

**Supplementary Fig. 1** Validation of CenSegNet centrosome segmentation. (page 2)

**Supplementary Fig. 2** Validation of inter-observer agreement for centrosome and epithelial annotations. (page 4)

**Supplementary Fig. 3** Centrosome segmentation performance of published deep learning models across imaging modalities. (page 5)

**Supplementary Fig. 4** Spatial uncoupling and inverse intracellular scaling between Stru CA and Num CA. (page 6)

**Supplementary Fig. 5** Centrinone-mediated PLK4 inhibition functionally uncouples structural and numerical centrosome abnormalities in hMECs. (page 7)

**Supplementary Fig. 6** Relationship between centrosome size and morphometric descriptors. (page 9)

**Supplementary Fig. 7** Patient age and tumour metrics across composite Stru CA and Num CA subgroups in edge and tumour regions. (page 10)

**Supplementary Fig. 8** Patient body composition and anthropometric parameters across composite Stru CA and Num CA subgroups in edge and tumour regions. (page 11)

**Supplementary Fig. 9** Numerical CA burden is associated with histological tumour grade in the edge region. (page 12)

**Supplementary Fig. 10** The burden of single-cell numerical CA is linked to histological tumour types. (page 13)

**Supplementary Fig. 11** CA patterns are hormone receptor-specific and influenced by the tumour microenvironment. (page 14)

**Supplementary Fig. 12** Kaplan–Meier analysis of overall survival (OS) and recurrence-free survival (RFS) stratified by Stru CA and Num CA status. (page 15)

**Supplementary Fig. 13** Multivariable Cox proportional hazards analyses of recurrence-free survival and overall survival. (page 17)

**Supplementary Fig. 14** Patient body composition and anthropometric parameters' associations with spatial dynamics of Stru CA and Num CA. (page 19)

**Supplementary Fig. 15** Spatial shifts in Stru CA and Num CA patterns are associated with distinct tumour characteristics. (page 20)

**Supplementary Fig. 16** Comparison of inference time across CPU and GPU configurations for the IHC dataset, IF dataset, and Epithelial dataset. (page 21)

**Supplementary Fig. 17** CenSegNet performance in appendix, colon, and kidney tissues. (page 22)

**Supplementary Fig. 18** Centrosome annotation across ground-truth datasets. (page 23)

### Supplementary Fig. 1

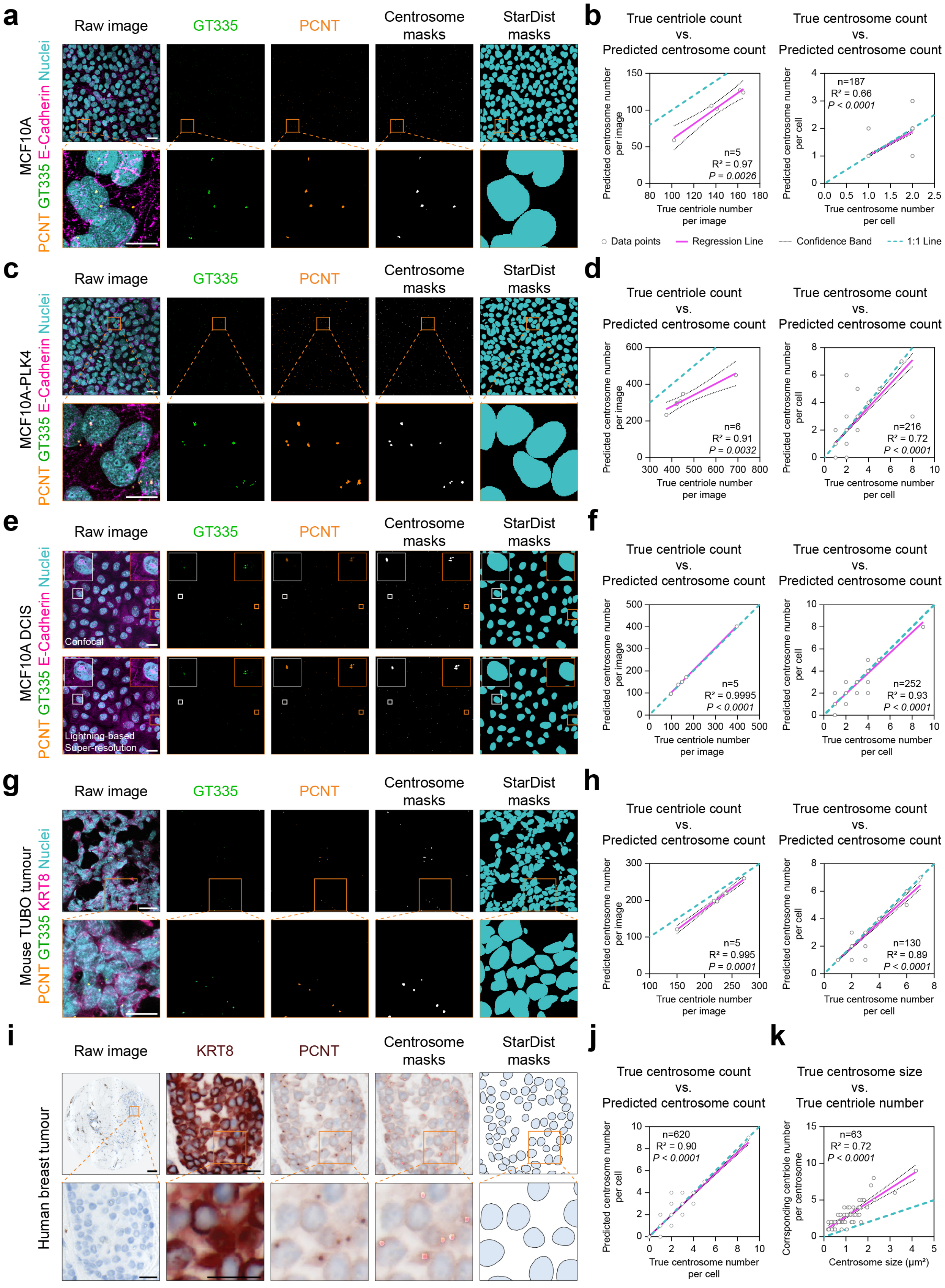


**Supplementary Fig. 1 Validation of CenSegNet centrosome segmentation. a, c, e, g** Representative immunofluorescence images of MCF10A, MCF10A-PLK4 cells, MCF10A DCIS cells (confocal and lightning-based super-resolution condition), and mouse TUBO tumour tissues (confocal) stained for pericentrin (PCNT: centrosome, orange), GT335 (centriole, green), and E cadherin or keratin 8 (KRT8, purple), and counterstained with DAPI (nuclei, teal). Corresponding centrosome and nuclear masks were generated by CenSegNet and StarDist. Scale bars, 20 μm. 2 **b, d, f, h** Left: true centriole number per image compared with predicted centrosome number by CenSegNet in MCF10A (n = 5 z-stack images, $R^2$ = 0.9666, **$P$ = 0.0026), MCF10A-PLK4 (n = 6 z-stack images, $R^2$ = 0.9085, **$P$ = 0.0032), MCF10A DCIS (n = 5 z-stack images, $R^2$ = 0.9995, ****$P$ < 0.0001), and mouse TUBO tissue (n = 5 z-stack images, $R^2$ = 0.9954, ****$P$ = 0.0001), Right: true centrosome number per cell compared with predicted centrosome number per cell in MCF10A (n=187 cells, $R^2$ = 0.6628, ****$P$ < 0.0001), MCF10A PLK4 (n=216 cells, $R^2$ = 0.7192, ****$P$ < 0.0001), MCF10A DCIS (n=252 cells, $R^2$ = 0.9309, ****$P$ < 0.0001) and mouse TUBO tissue (n=130 cells, $R^2$ = 0.8881, ****$P$ < 0.0001). **i** Representative immunohistochemistry image of human breast tumour tissue stained for KRT8 and PCNT, and counterstained with haematoxylin (nuclei), with centrosomes and nuclei segmented by CenSegNet and StarDist. **j** Correlation between true and predicted centrosome number per cell (n=620 cells, $R^2$ = 0.9005, ****$P$ < 0.0001). **k** Correlation between centrosome size and corresponding centriole number in MCF10A-PLK4 cells (n = 63 centrosomes, $R^2$ = 0.7235, ****$P$ < 0.0001). Data are presented as individual data points. Source data are provided as a Source Data file.

#### Supplementary Fig. 2

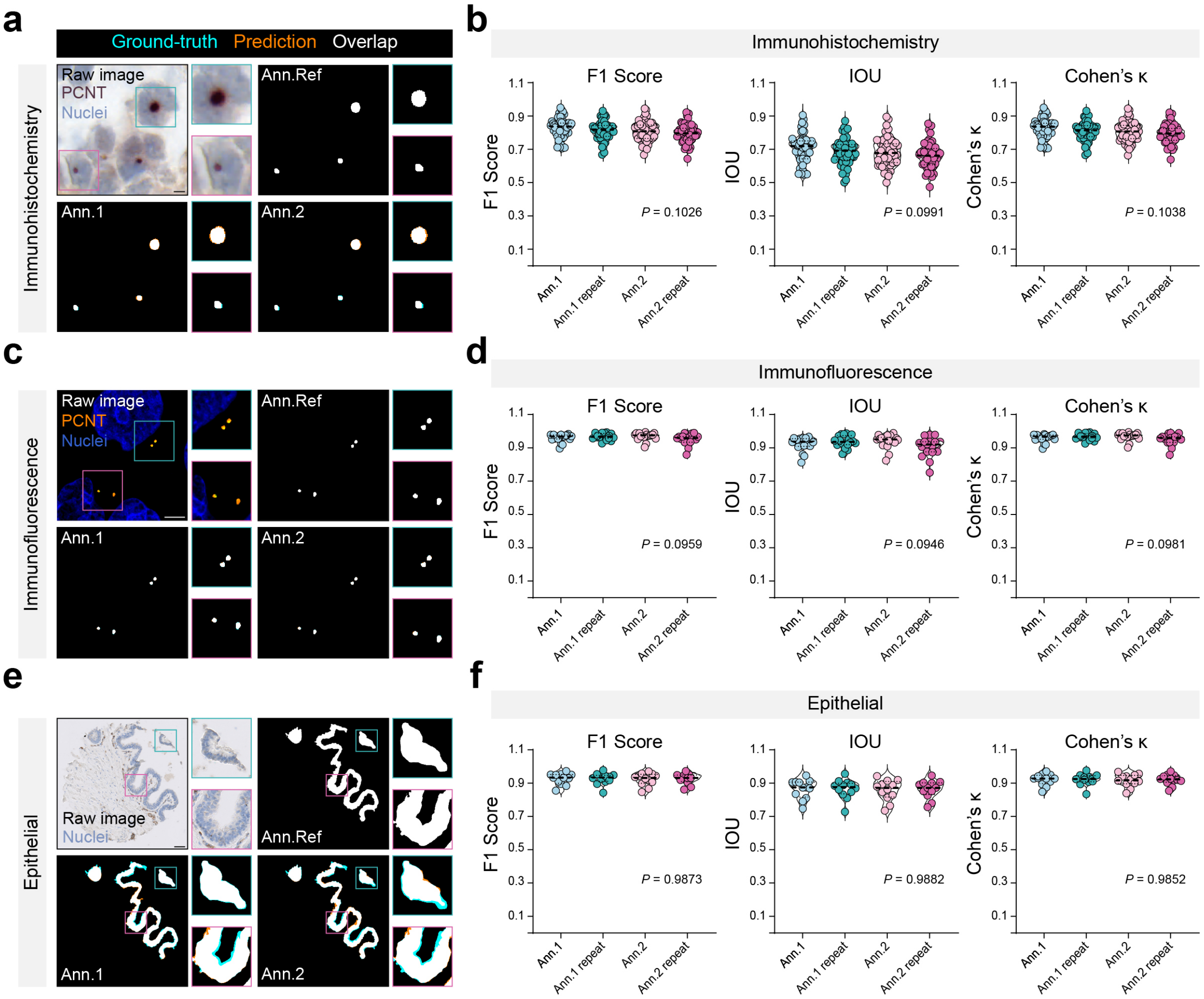


**Supplementary Fig. 2 Validation of inter-observer agreement for centrosome and epithelial annotations. a, c, e** Representative immunohistochemistry image of human breast tumour tissue stained for PCNT (**a**) and counterstained with haematoxylin (nuclei) (**a, e**) and immunofluorescence images (confocal) stained for PCNT and counterstained with DAPI (nuclei) (**c**) used for centrosome (**a, c**) and epithelial (**e**) annotation. Shown are raw images alongside annotations from the reference annotator (Ann. Ref), annotator 1 (Ann. 1) and annotator 2 (Ann. 2). Overlay maps depict agreement between annotations: teal, regions present only in Ann. Ref; orange, regions present only in Ann.1 or Ann.2; white, overlapping regions. Scale bars, 5 μm (**a, c**) and 50 μm (**e**). **b, d, f** Quantification of inter-observer agreement using F1 score, intersection over union (IoU) and Cohen's κ for centrosome (**b, d**) and epithelial (**f**) annotations in immunohistochemistry (**b, f**) and immunofluorescence (**d**). Ann.1 and Ann.2 were each evaluated in initial and repeat annotation rounds. Each point represents an individual 256 × 256 image patch in **b, d** and a whole tissue core in **f**; centre lines indicate mean values. Statistical analysis was performed using one-way ANOVA with Tukey's post hoc test; no significant differences were detected (exact *P* values shown). Data are presented as individual data points. Source data are provided as a Source Data file.

#### Supplementary Fig. 3

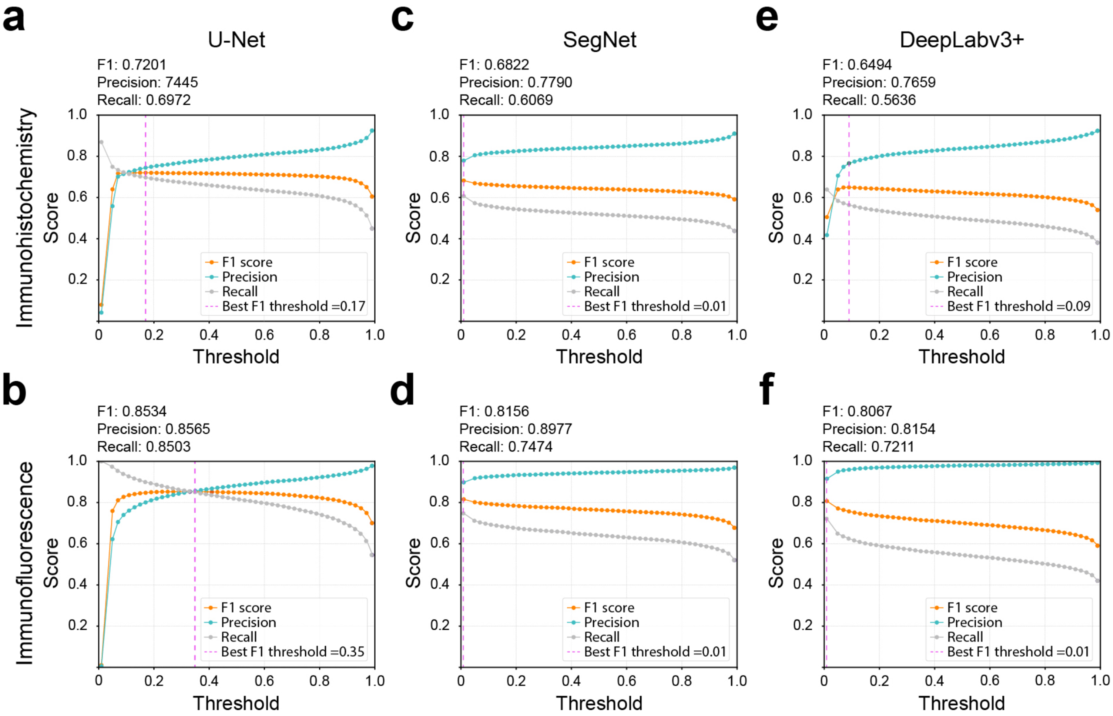


**Supplementary Fig. 3 Centrosome segmentation performance of published deep learning models across imaging modalities. a-f** Precision, recall, and F1 score for threshold optimisation of U-Net, SegNet, DeepLabv3+ models, evaluated on immunohistochemistry (**a, c, e**) and immunofluorescence (**b, d, f**) images. The plots show the relationship between threshold values and model performance (orange: F1 score; blue: precision; grey: recall; dashed magenta line: optimal threshold) against different thresholds (*x*-axis). The F1 score, precision, and recall values at the best threshold are reported in the top left of each plot. Each model is evaluated using the metrics at the best threshold: **a** U-Net (F1 score = 0.7201, precision = 0.7445, recall = 0.6972), **b** SegNet (F1 score = 0.6822, precision = 0.7790, recall = 0.6069), **c** DeepLabv3+ (F1 score = 0.6494, precision = 0.7659, recall = 0.5636), **d** U-Net (F1 score = 0.8534, precision = 0.8565, recall = 0.8503), **e** SegNet (F1 score = 0.8156, precision = 0.8977, recall = 0.7474), and **f** DeepLabv3+ (F1 score = 0.8067, precision = 0.8154, recall = 0.7211). The results show model performance variation depending on imaging modality and the threshold applied.

### Supplementary Fig. 4

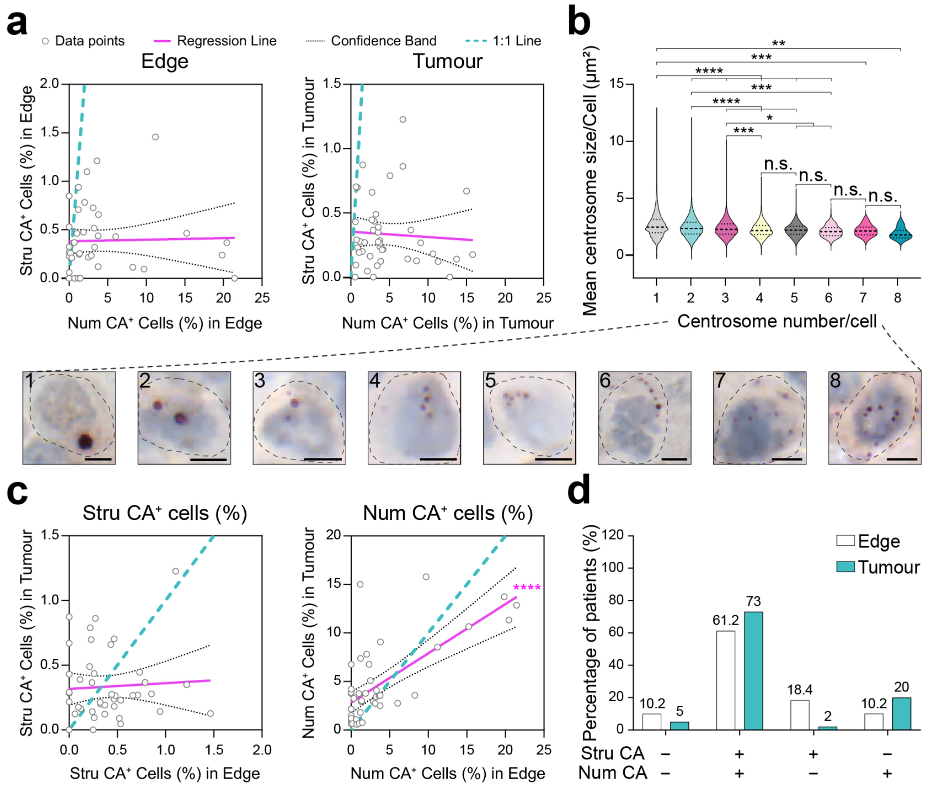


**Supplementary Fig. 4 Spatial uncoupling and inverse intracellular scaling between Stru CA and Num CA. a** Left: Pearson correlation between the percentage of cells with Stru CA and Num CA in the edge region. Right: Pearson correlation between the percentage of cells with Stru CA and Num CA in the tumour region. Grey circles represent individual patients, purple line represents regression line, Black dot line represents confidence band, and teal dot line represents 1:1 line. Two-tailed Pearson correlation test followed by simple linear regression for visualisation, left: $R^2 = 0.0007$, $P = 0.8663$; right: $R^2 = 0.0041$, $P = 0.6843$. **b** Relationship between centrosome number and centrosome size per cell and corresponding representative immunohistochemistry images of human breast tumour tissues stained for PCNT (centrosome) and counterstained with haematoxylin (nuclei). *x*-axis indicates the centrosome number per cell; *y*-axis indicates the mean centrosome size per cell. One-way ANOVA (****$P < 0.0001$) with Tukey's test, *$P$ (3 vs 5) = 0.0178; *$P$ (3 vs 6) = 0.0304; ** $P = 0.0091$; top: ***$P = 0.0001$, middle: ***$P = 0.0002$, bottom: ***$P = 0.0004$; ****$P < 0.0001$. n.s. (not significant). Data are presented as violin plots showing the distribution of values; dashed lines indicate median and interquartile ranges. Each image is labelled with the number of centrosomes per cell; black dotted lines outline cell boundaries. Scale bars, 5 µm. **c** Left: Pearson correlation of the percentage of cells with Stru CA between edge and tumour regions. Right: Pearson correlation of the percentage of cells with Num CA between edge and tumour regions. Grey circles represent individual patients, purple line represents regression line, Black dot line represents confidence band, and teal dot line represents 1:1 line. Two-tailed Pearson correlation test, followed by simple linear regression for visualisation, left: $R^2 = 0.0033$, $P = 0.7137$; right: $R^2 = 0.4856$, ****$P < 0.0001$. **d** Patients were classified by composite CA burden into $Stru^-Num^-$ (Edge: n = 10 patients; Tumour: n = 5 patients), $Stru^+Num^+$ (Edge: n = 30 patients; Tumour: n = 73 patients), $Stru^+Num^-$ (Edge: n = 9 patients; Tumour: n = 2 patients), $Stru^-Num^+$ (Edge: n = 5 patients; Tumour: n = 20 patients) groups. Histograms show the percentage of patients in each group in edge and tumour regions. Source data are provided as a Source Data file.

#### Supplementary Fig. 5

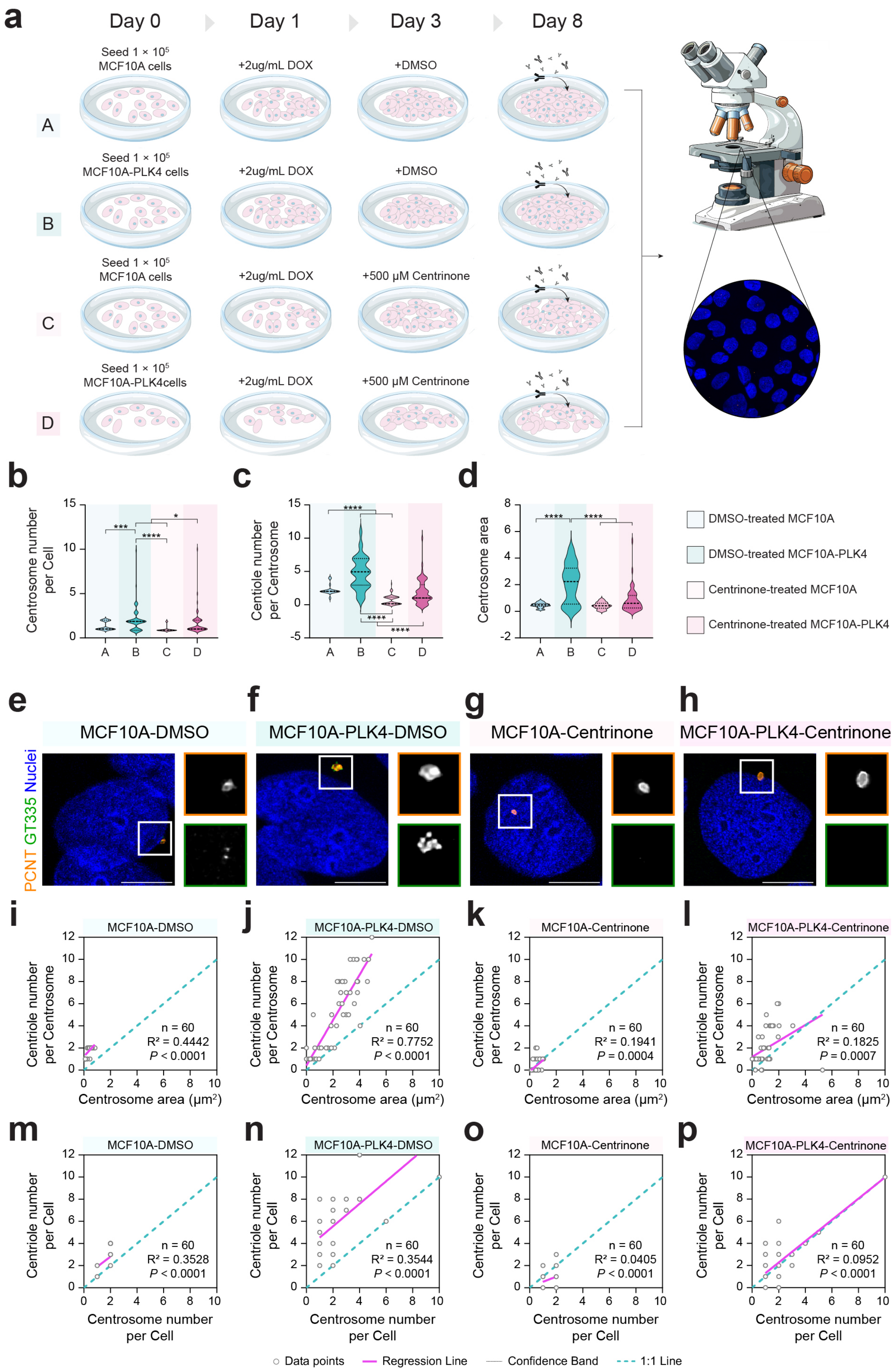


**Supplementary Fig. 5 Centrinone-mediated PLK4 inhibition functionally uncouples structural and numerical centrosome abnormalities in hMECs. a** Schematic of the experimental design. MCF10A-PLK4 cells were treated with 2 μg ml⁻¹ doxycycline (DOX) for 48

h to induce centriole amplification. Cells were then treated with the PLK4-selective inhibitor Centrinone for 5 days. Four experimental groups (A–D) were defined based on cell line and treatment conditions, with MCF10A cells serving as controls. At day 8, cells were fixed and analysed by confocal microscopy following immunostaining for centrosomal and centriolar markers. **b, c, d** Quantification of centrosome number per cell (b), centriole number per centrosome (c), and centrosome area (d) across groups A–D. Data were analysed by one-way ANOVA (****$P < 0.0001$) with Tukey's multiple comparisons test, [*$P$ (B vs. D) = 0.0199; *$P$ (C vs. D) = 0.0329; ***$P = 0.0001$, ****$P < 0.0001$]. **e-h** Representative lightning-based super-resolution images of MCF10A and MCF10A-PLK4 cells stained for pericentrin (PCNT; centrosomes, orange) and GT335 (centrioles, green), with nuclei counterstained using DAPI (blue). Insets show magnified views of centrosomes (orange boxes) and centrioles (green boxes). Scale bars, 5 µm. **i-l** Correlation between centrosome area and centriole number per centrosome across groups A-D. **m-p** Correlation between centrosome number per cell and centriole number per cell across groups A-D. Grey circles represent individual cells, purple line represents regression line, Black dot line represents confidence band, and teal dot line represents 1:1 line. Two-tailed Pearson correlation test followed by simple linear regression for visualisation. Data are presented as individual data points. Source data are provided as a Source Data file.

### Supplementary Fig. 6

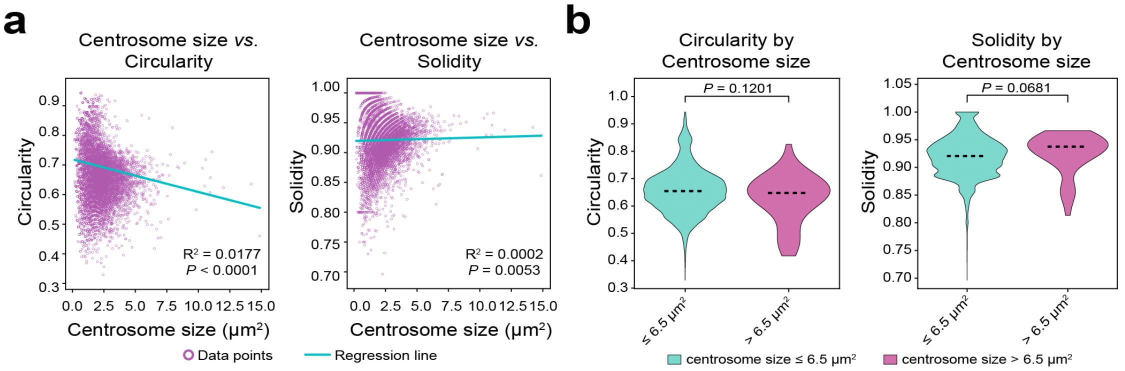


**Supplementary Fig. 6 Relationship between centrosome size and morphometric descriptors. a** Scatter plots showing the relationship between centrosome size and circularity and solidity in the annotation immunohistochemistry dataset (n = 15185). Each purple circle represents an individual centrosome. Blue lines denote fitted regression lines. **b** Violin plots comparing circularity and solidity between normal (⩽ 6.5 μm$^2$; n = 15145) and enlarged (> 6.5 μm$^2$; n = 40) centrosomes. Dashed lines indicate medians. For **a**, two-tailed Pearson correlation tests were performed, with simple linear regression shown for visualisation; $P$ and $R^2$ values are indicated in the corresponding panels. For **b**, comparisons were performed using two-sided Mann–Whitney U tests; $P$ values are indicated in the corresponding panels. Source data are provided as a Source Data file.

### Supplementary Fig. 7

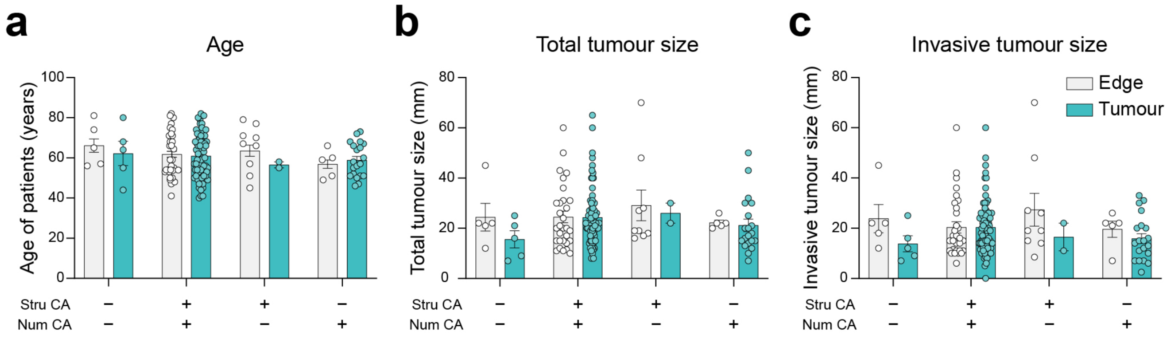


**Supplementary Fig. 7 Patient age and tumour metrics across composite Stru CA and Num CA subgroups in edge and tumour regions. a–c** Comparative analysis of patient age, total tumour size, and histological tumour size across Stru$^{-}$Num$^{-}$, Stru$^{+}$Num$^{+}$, Stru$^{+}$Num$^{-}$, Stru$^{-}$Num$^{+}$ CA groups in Edge and Tumour regions. Two-way ANOVA (Age, $P$ = 0.3606; Total tumour size, $P$ = 0.4922; Histological tumour size, $P$ = 0.7985) with Sidak's test: Age (Edge *versus* Tumour: $P$ = 0.3825; Stru$^{-}$Num$^{-}$–Stru$^{-}$Num$^{+}$ CA groups: $P$ = 0.3606*)*; Total tumour size (Edge *versus* Tumour: $P$ = 0.3391; Stru$^{-}$Num$^{-}$–Stru$^{-}$Num$^{+}$ CA groups: $P$ = 0.4922*)*; Histological tumour size (Edge *versus* Tumour: $P$ = 0.0613; Stru$^{-}$Num$^{-}$–Stru$^{-}$Num$^{+}$ CA groups: $P$ = 0.7985*)*, absence of asterisks indicates no statistical significance. Data are presented as individual data points and mean ± s.e.m. Source data are provided as a Source Data file.

### Supplementary Fig. 8

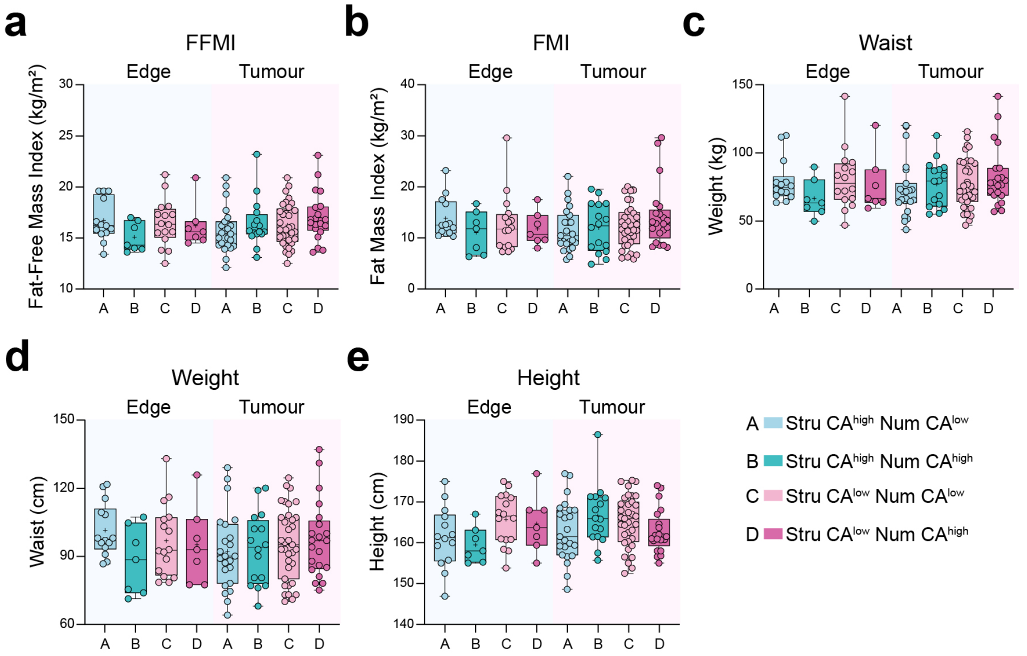


**Supplementary Fig. 8 Patient body composition and anthropometric parameters across composite Stru CA and Num CA subgroups in edge and tumour regions. a–e.** Patients classified by composite CA burden into $Stru^{high}Num^{low}$ (A), $Stru^{high}Num^{high}$ (B), $Stru^{low}Num^{low}$ (C), and $Stru^{low}Num^{high}$ (D) groups in Edge (A: n = 14 patients; B: n = 7 patients; C: n = 16 patients; D: n = 7 patients) and Tumour (A: n = 20 patients; B: n = 16 patients; C: n = 35 patients; D: n = 23 patients) regions. Comparative analysis of patient characteristics, including fat-free mass index (FFMI), fat mass index (FMI), height, waist circumference, and weight across A–D groups in edge and tumour regions. Kruskal-Wallis H Test with Dunn's multiple comparison test was performed in edge and tumour regions separately. Edge (FFMI: $P$ = 0.4356; FMI: $P$ = 0.6005; Waist: $P$ = 0.2455; Weight: $P$ = 0.2474; Height: $P$ = 0.1298), Tumour (FFMI: $P$ = 0.1927; FMI: $P$ = 0.5277; Waist: $P$ = 0.9982; Weight: $P$ = 0.3680; Height: $P$ = 0.2125). Data are presented as individual data points and as box and whiskers plots showing the distribution of values, median and quartiles. Source data are provided as a Source Data file.

### Supplementary Fig. 9

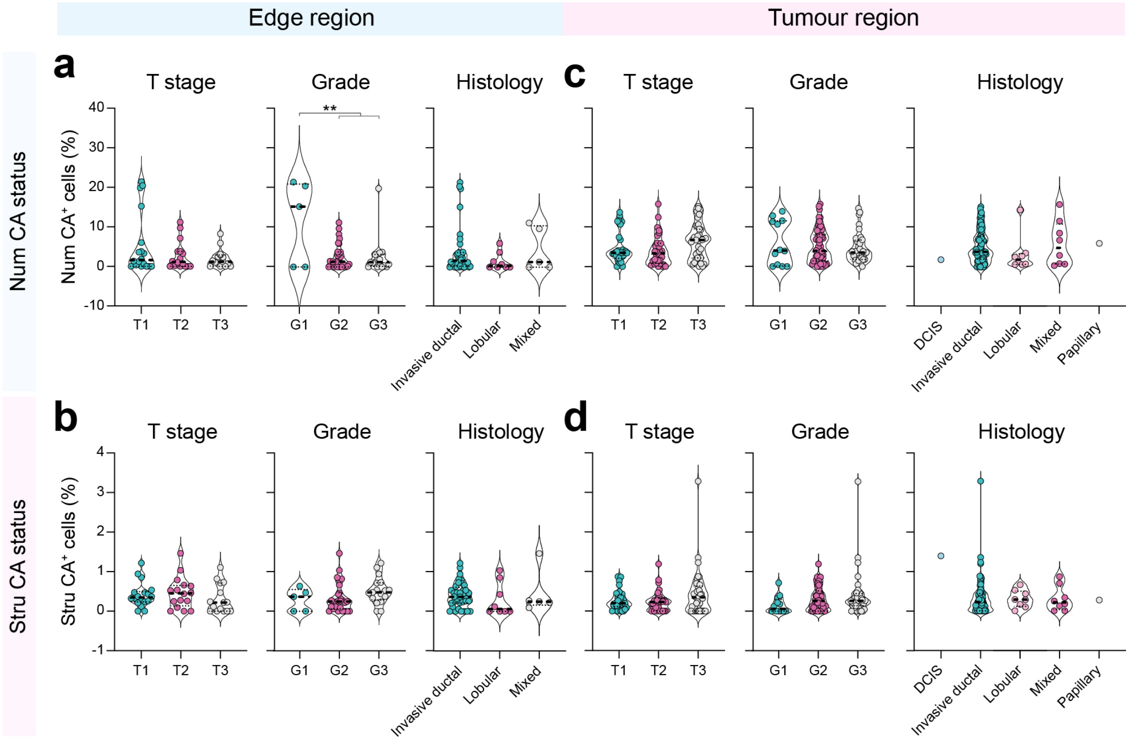


**Supplementary Fig. 9 Numerical CA burden is associated with histological tumour grade in the edge region. a–d** Comparative analysis of the percentage of cells with Stru CA and Num CA stratified by T stage, histological tumour grade, and histological tumour type in edge and tumour regions. One-way ANOVA [Num CA status in Edge (T stage, $P$ = 0.1482; Grade, $^{**}P$ = 0.0017; Histological type, $P$ = 0.5065); Num CA status in Tumour (T stage, $P$ = 0.1386; Grade, $P$ = 0.1386; Histological type, $P$ = 0.6568); Stru CA status in Edge (T stage, $P$ = 0.5021 ; Grade, $P$ = 0.2223; Histological type, $P$ = 0.6982); Stru CA status in Tumour (T stage, $P$ = 0.1626; Grade, $P$ = 0.2206; Histological type, $P$ = 0.2051)] with Tukey's test, Grade, $^{**}P$ (G1 vs G2) = 0.0017, $^{**}P$ (G1 vs G3) = 0.0022. Data are presented as individual data points and mean ± s.e.m. Source data are provided as a Source Data file.

### Supplementary Fig. 10

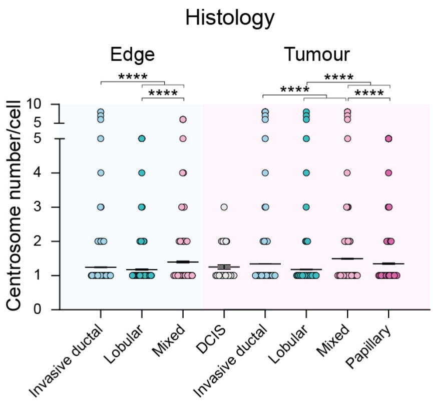


**Supplementary Fig. 10 The burden of single-cell numerical CA is linked to histological tumour types.** Comparative analysis of centrosome number per cell across different Histological tumour types in edge and tumour regions. Kruskal-Wallis H Test with Dunn's multiple comparison test was performed in edge and tumour regions separately. Edge: ****$P < 0.0001$; Tumour: ****$P < 0.0001$. Data are presented as individual data points and mean ± s.e.m. Source data are provided as a Source Data file.

### Supplementary Fig. 11

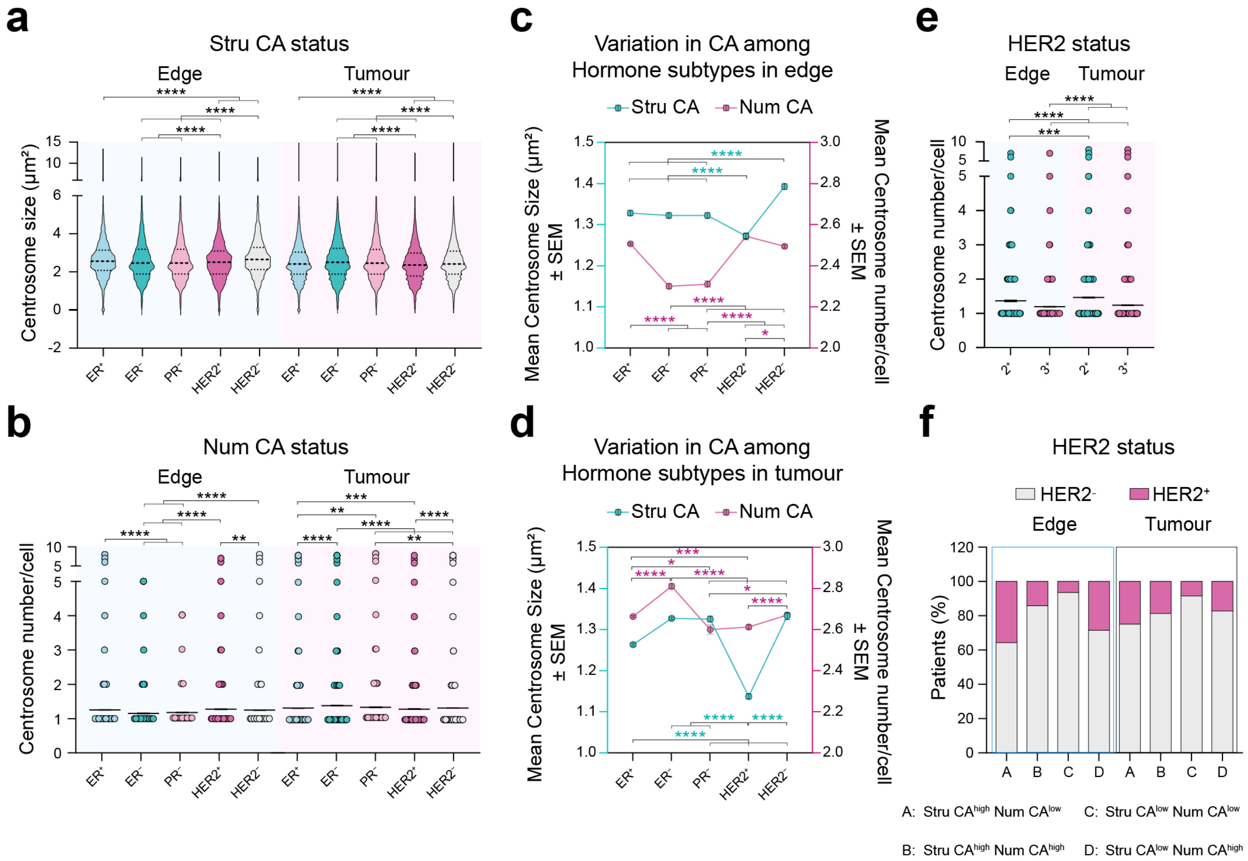


**Supplementary Fig. 11 CA patterns are hormone receptor-specific and influenced by the tumour microenvironment. a** Comparative analysis of centrosome size across different hormone status in edge and tumour regions. One-way ANOVA (Edge: ****$P$ < 0.0001; Tumour: ****$P$ < 0.0001) with Tukey's test was performed, Edge: ****$P$ < 0.0001; Tumour: ****$P$ < 0.0001. Data are presented as violin plots showing the distribution of values; dashed lines indicate median and interquartile ranges. **b** Comparative analysis of centrosome number per cell across different hormone status in edge and tumour regions. Kruskal-Wallis H Test with Dunn's multiple comparison test was performed, Edge: **$P$ = 0.0077, ****$P$ < 0.0001; Tumour: (top)**$P$ = 0.0090, (bottom)**$P$ = 0.0035, ***$P$ = 0.0004, ****$P$ < 0.0001. Data are presented as individual data points and mean ± s.e.m. **c, d** Variation of centrosome size number per cell across hormone subtypes in edge and tumour regions. One-way ANOVA (Edge, ****$P$ < 0.0001; Tumour, ****$P$ < 0.0001) with Tukey's test, Edge (Stru CA: ****$P$ < 0.0001; Num CA: *$P$ = 0.0316, ****$P$ < 0.0001), Tumour [Stru CA: ****$P$ < 0.0001; Num CA: (top) *$P$ = 0.0471, (bottom) *$P$ = 0.0201, ***$P$ = 0.0004, ****$P$ < 0.0001]. Data are presented as mean ± s.e.m. **e** Comparative analysis of centrosome number per cell across different HER2 status in edge and tumour regions. HER2 status was evaluated by immunohistochemistry. Kruskal-Wallis H Test with Dunn's multiple comparison test, ***$P$ = 0.0002, **** $P$ < 0.0001. Data are presented as individual data points and mean ± s.e.m. **f** Patients classified by composite CA burden into $\text{Stru}^{high}\text{Num}^{low}$ (A), $\text{Stru}^{high}\text{Num}^{high}$ (B), $\text{Stru}^{low}\text{Num}^{low}$ (C), and $\text{Stru}^{low}\text{Num}^{high}$ (D) groups in Edge (A: n = 14 patients; B: n = 7 patients; C: n = 16 patients; D: n = 7 patients) and Tumour (A: n = 20 patients; B: n = 16 patients; C: n = 35 patients; D: n = 23 patients) regions. Histograms show the percentage of patients with HER2 status across composite CA groups A–D in edge and tumour regions. Fisher exact test, absence of asterisks indicates no statistically significant correlation. Source data are provided as Source Data file.

### Supplementary Fig. 12

**a**

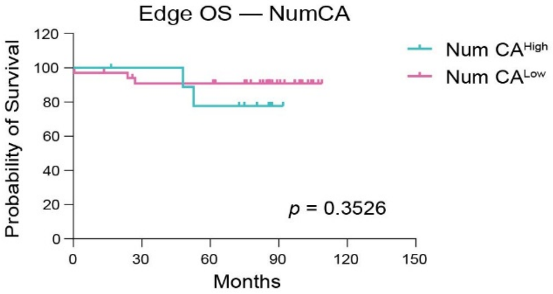


Number at risk (number censored)

| | | 30 | 60 | 90 | 120 | 150 |
|---|---|---|---|---|---|---|
| Num CA$^{High}$ | 10 (0) | 9 (1) | 7 (1) | 1 (7) | 0 (8) | 0 (8) |
| Num CA$^{Low}$ | 34 (0) | 29 (2) | 29 (2) | 12 (19) | 0 (31) | 0 (31) |

**b**

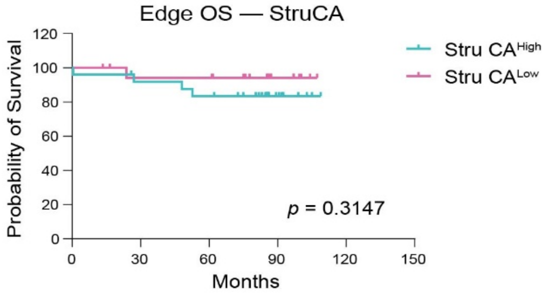


Number at risk (number censored)

| | | 30 | 60 | 90 | 120 | 150 |
|---|---|---|---|---|---|---|
| Stru CA$^{High}$ | 25 (0) | 22 (1) | 20 (1) | 7 (14) | 0 (21) | 0 (21) |
| Stru CA$^{Low}$ | 19 (0) | 16 (2) | 16 (2) | 6 (12) | 0 (18) | 0 (18) |

**c**

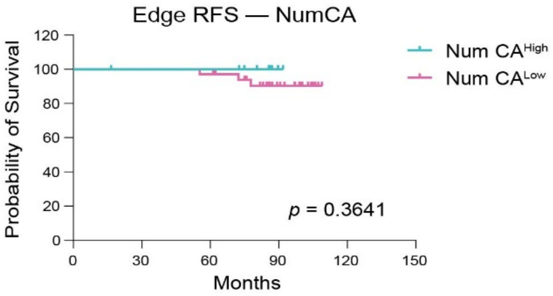


Number at risk (number censored)

| | | 30 | 60 | 90 | 120 | 150 |
|---|---|---|---|---|---|---|
| Num CA$^{High}$ | 10 (0) | 9 (1) | 9 (1) | 1 (9) | 0 (10) | 0 (10) |
| Num CA$^{Low}$ | 34 (0) | 34 (0) | 33 (0) | 14 (17) | 0 (31) | 0 (31) |

**d**

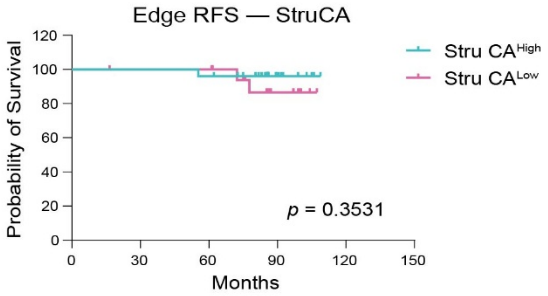


Number at risk (number censored)

| | | 30 | 60 | 90 | 120 | 150 |
|---|---|---|---|---|---|---|
| Stru CA$^{High}$ | 25 (0) | 25 (0) | 24 (0) | 8 (16) | 0 (24) | 0 (24) |
| Stru CA$^{Low}$ | 19 (0) | 18 (1) | 18 (1) | 7 (10) | 0 (17) | 0 (17) |

**e**

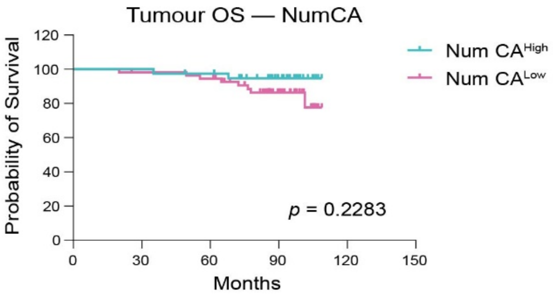


Number at risk (number censored)

| | | 30 | 60 | 90 | 120 | 150 |
|---|---|---|---|---|---|---|
| Num CA$^{High}$ | 39 (0) | 39 (0) | 37 (1) | 19 (18) | 0 (37) | 0 (37) |
| Num CA$^{Low}$ | 56 (0) | 54 (1) | 52 (1) | 25 (24) | 0 (48) | 0 (48) |

**f**

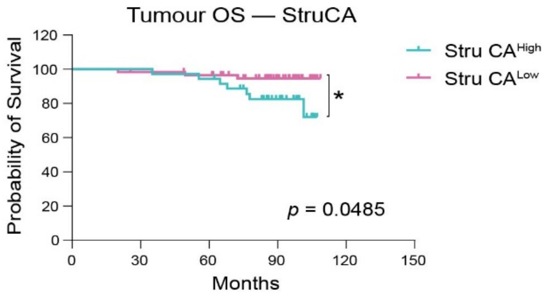


Number at risk (number censored)

| | | 30 | 60 | 90 | 120 | 150 |
|---|---|---|---|---|---|---|
| Stru CA$^{High}$ | 36 (0) | 36 (0) | 34(0) | 18 (12) | 0 (29) | 0 (29) |
| Stru CA$^{Low}$ | 59 (0) | 57 (1) | 55 (2) | 26 (30) | 0 (56) | 0 (56) |

**g**

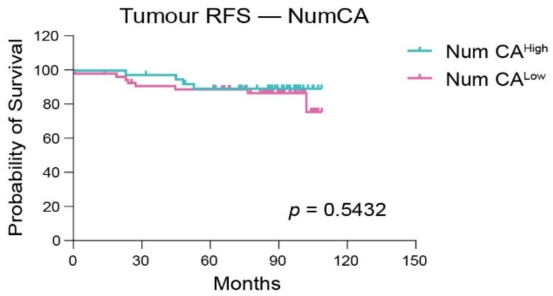


Number at risk (number censored)

| | | 30 | 60 | 90 | 120 | 150 |
|---|---|---|---|---|---|---|
| Num CA$^{High}$ | 39 (0) | 38 (0) | 33 (2) | 18 (17) | 0 (35) | 0 (35) |
| Num CA$^{Low}$ | 56 (0) | 49 (2) | 48 (2) | 23 (26) | 0 (48) | 0 (48) |

**h**

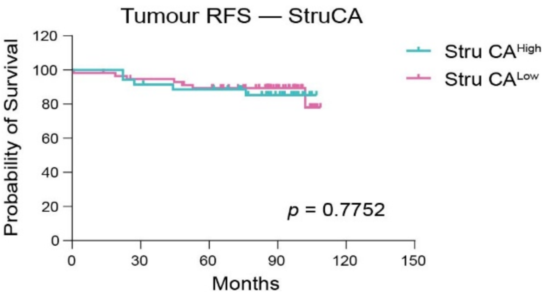


Number at risk (number censored)

| | | 30 | 60 | 90 | 120 | 150 |
|---|---|---|---|---|---|---|
| Stru CA$^{High}$ | 36 (0) | 33 (0) | 31 (1) | 16 (15) | 0 (31) | 0 (31) |
| Stru CA$^{Low}$ | 59 (0) | 54 (2) | 50 (3) | 25 (28) | 0 (52) | 0 (52) |

**Supplementary Fig. 12 Kaplan–Meier analysis of overall survival (OS) and recurrence-free survival (RFS) stratified by Stru CA and Num CA status.** Cyan curves represent cases with high Stru CA or Num CA levels, whereas magenta curves represent cases with low levels. **a–d** correspond to analyses based on CA status in the edge region, and Figures **e–h** corresponds to analyses based on CA status in the tumour region. Tick marks indicate censored cases, and numbers at risk are shown below each panel. Patients were stratified according to composite CA burden into Stru $CA^{High}$ and Stru $CA^{Low}$ groups or Num $CA^{High}$ and Num $CA^{Low}$ groups in the edge and tumour regions, respectively. **a, c** Num $CA^{High}$, n = 10 patients; Num $CA^{Low}$, n = 34 patients. **b, d** Stru $CA^{High}$, n = 25 patients; Stru $CA^{Low}$, n = 19 patients. **e** Num $CA^{High}$, n = 39 patients; Num $CA^{Low}$, n = 56 patients. **f** Stru $CA^{High}$, n = 36 patients; Stru $CA^{Low}$, n = 59 patients. **g** Num $CA^{High}$, n = 39 patients; Num $CA^{Low}$, n = 56 patients. **h** Stru $CA^{High}$, n = 36 patients; Stru $CA^{Low}$, n = 59 patients. Survival curves were estimated using the Kaplan–Meier method, and differences between groups were assessed using the log-rank test. *P* values were calculated using the log-rank test and are displayed in the corresponding figures.

### Supplementary Fig. 13

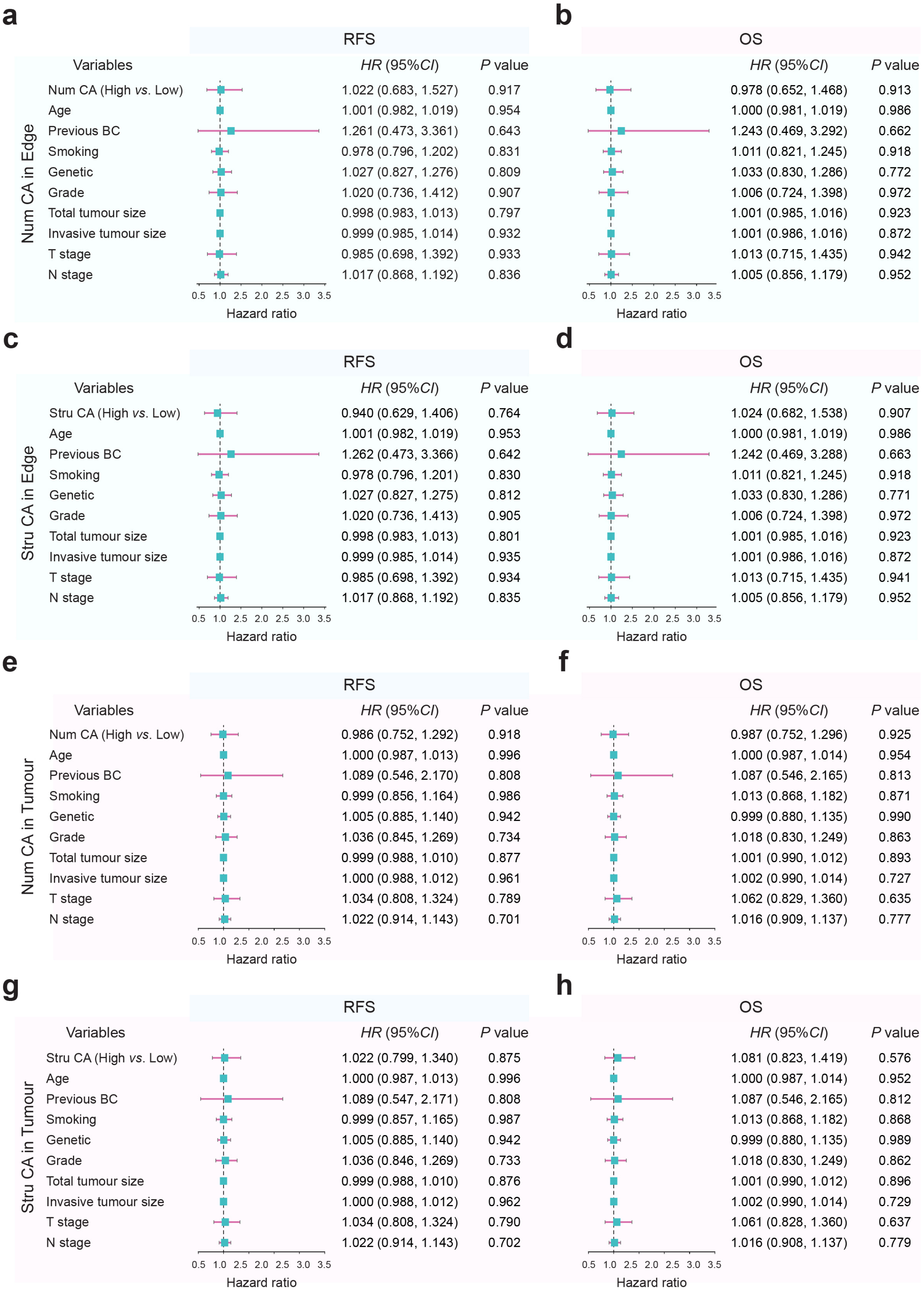


**Supplementary Fig. 13 Multivariable Cox proportional hazards analyses of recurrence-free survival and overall survival.** Forest plots showing hazard ratios (HRs) and 95% confidence intervals (CIs) derived from multivariable Cox proportional hazards models for recurrence-free survival (RFS; left) and overall survival (OS; right). Analyses were performed for CA measurements in different anatomical regions: **a, b** Num CA in edge region, n = 44 patients; **c, d** Stru CA in edge region, n = 44 patients; **e, f** Num CA in tumour region, n = 95 patients; and **g, h** Stru CA in tumour region, n = 95 patients. In each model, CA was included as a dichotomized variable (High *vs*. Low based on medium value). All models were adjusted for established clinicopathological factors, including age, previous breast cancer history, smoking status, genetic status, tumour grade, total tumour size, invasive tumour size, tumour (T) stage, and nodal (N) status. Squares indicate HR estimates, and horizontal lines represent 95% CIs. The vertical dashed line denotes HR = 1. *P* values were calculated using two-sided Wald tests. RFS, recurrence-free survival; OS, overall survival; HR, hazard ratio; CI, confidence interval.

#### Supplementary Fig. 14

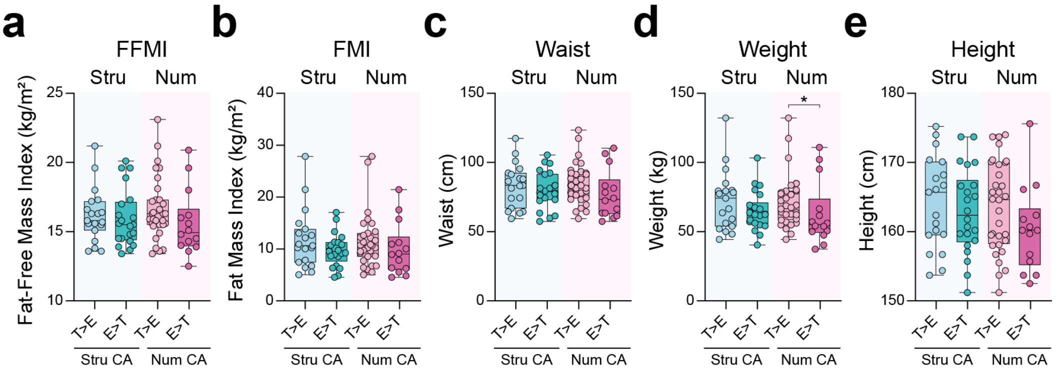


**Supplementary Fig. 14 Patient body composition and anthropometric parameters' associations with spatial dynamics of Stru CA and Num CA. a–e** Patients were stratified by relative CA burden between edge and tumour regions into four groups: Stru $CA^{T>E}$ (Stru CA enriched in the tumour region), Stru $CA^{E>T}$ (Stru CA enriched in the edge region), Num $CA^{T>E}$ (Num CA enriched in the tumour region), Num $CA^{E>T}$ (Num CA enriched in the edge region). Histograms show patient FFMI, FMI, waist, weight, and height across groups. Mann-Whitney U test, FFMI (Stru, $P$ = 0.4980; Num $P$ = 0.0969); FMI (Stru, $P$ = 0.2389; Num $P$ = 0.2255); Waist (Stru, $P$ = 0.4720; Num $P$ = 0.0720); Weight (Stru, $P$ = 0.3746; Num *$P$ = 0.0339); Height (Stru, $P$ = 0.5115; Num $P$ = 0.1661). Data are presented as individual data points and as box and whiskers plots showing the distribution of values, median and quartiles. Source data are provided as Source Data file.

### Supplementary Fig. 15

**a**

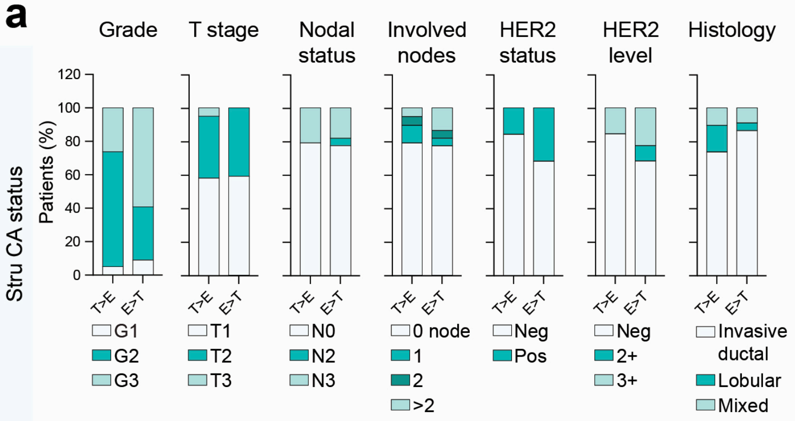


**b**

characteristics with >10% differences

| Tumour characteristics | T>E | E>T |
|---|---|---|
| Quickly growing (Grade 3) (%) | 26.3 | 59.1 |
| HER2 status Positive (%) | 15.8 | 31.8 |
| Tumour size>2mm (%) | 42.1 | 40.9 |
| Nodal status (N3) (%) | 21.8 | 18.2 |
| Histology-Mixed (%) | 10.6 | 9.1 |
| Histology-Lobular (%) | 15.8 | 4.6 |

**c**

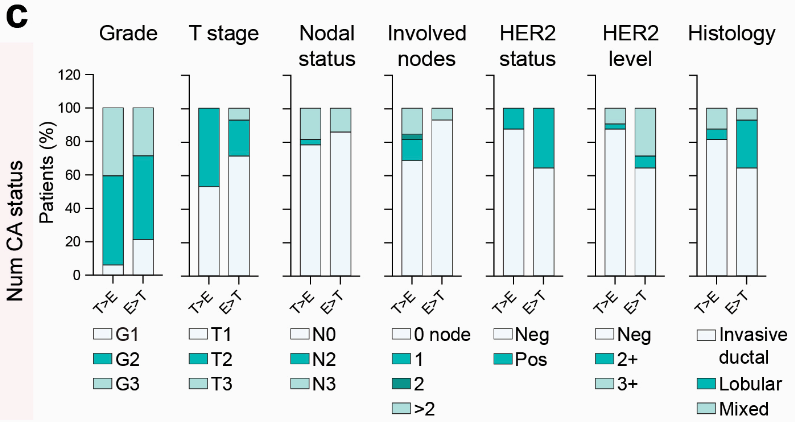


**d**

characteristics with >10% differences

| Tumour characteristics | T>E | E>T |
|---|---|---|
| Quickly growing (Grade 3) (%) | 40.6 | 28.6 |
| HER2 status Positive (%) | 12.5 | 35.7 |
| Tumour size>2mm (%) | 46.9 | 28.5 |
| Nodal status (N3) (%) | 18.8 | 14.3 |
| Histology-Mixed (%) | 12.5 | 7.1 |
| Histology-Lobular (%) | 6.3 | 28.6 |

**Supplementary Fig. 15 Spatial shifts in Stru CA and Num CA patterns are associated with distinct tumour characteristics. a, c** Percentage of patients with different tumour characteristics (histological tumour grade, T stage, nodal status, involved nodes number, HER2 status, HER2 expression level, and histological tumour types) across the Stru CA$^{T>E}$, Stru CA$^{E>T}$, Num CA$^{T>E}$, and Num CA$^{E>T}$ groups. Fisher exact test, absence of asterisks indicates no statistically significant correlation. **a** Grade, *P* = 0.0510; T stage, *P* = 0.7423; Nodal status, *P* >0.9999; Involved nodes, *P* = 0.7884; HER2 status, *P* = 0.2919; HER2 level, *P* = 0.5303; Histological type, *P* = 0.6185. **c** Grade, *P* = 0.3205; T stage, *P* = 0.0899; Nodal status, *P* >0.9999; Involved nodes, *P* = 0.4796; HER2 status, *P* = 0.1060; HER2 level, *P* = 0.1988; Histological type, *P* = 0.1311. **b, d** Tables summarising the percentage of tumours exhibiting specific characteristics [Quickly growing (Grade 3), HER2 status Positive, Tumour size > 2 mm, Nodal status (N3), Histology-Mixed, and Histology-Lobular] categorised by a spatial shift in CA patterns. Highlighted purple cells indicate characteristics with higher than 10% differences in the percentages between the T>E and E>T groups. Source data are provided as Source Data file.

### Supplementary Fig. 16

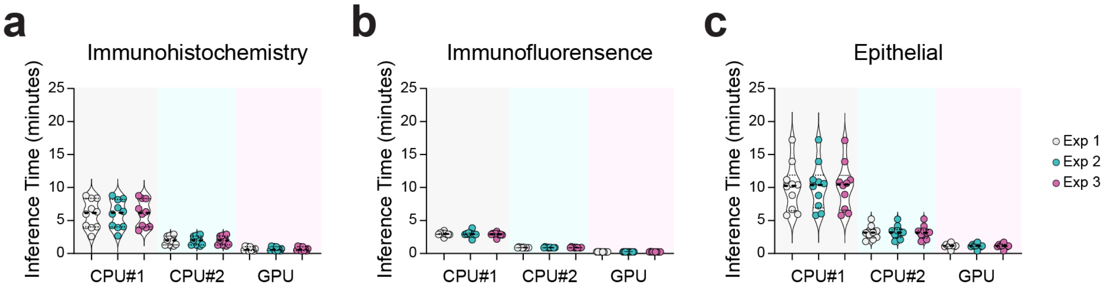


**Supplementary Fig. 16 Comparison of inference time across CPU and GPU configurations for the IHC dataset (a), IF dataset (b), and Epithelial dataset (c).** Dots represent individual runs from three experiments (Exp 1–3), with colours indicating experiment identity. Summary bars show mean ± SEM. The immunofluorescence dataset consisted of 10 images, each with a resolution of 1024 × 1024 pixels. The immunohistochemistry and epithelial (haematoxylin-stained) datasets each consisted of 10 TMA core images, with image resolutions ranging from 3370 × 4487 to 6630 × 5941 pixels. GPU consistently reduced computation time relative to CPU#1 and CPU#2 across all datasets. CPU#1 is an office laptop with 11th Gen Intel(R) Core (TM) i5-1145G7 @ 2.60GHz (1.50 GHz) processor with 8 GB RAM. CPU#2 is a server with AMD EPYC 9334 32-Core Processor with 128 GB RAM. GPU is an NVIDIA Tesla T4 approximately 16 GB VRAM, which is the standard "free-tier" GPU provided by Google Colab. Data are presented as individual data points and mean ± s.e.m. The pixel dimensions of each image used for benchmarking are provided in the Source Data to facilitate interpretation of the reported inference times. Source data are provided as a Source Data file.

### Supplementary Fig. 17

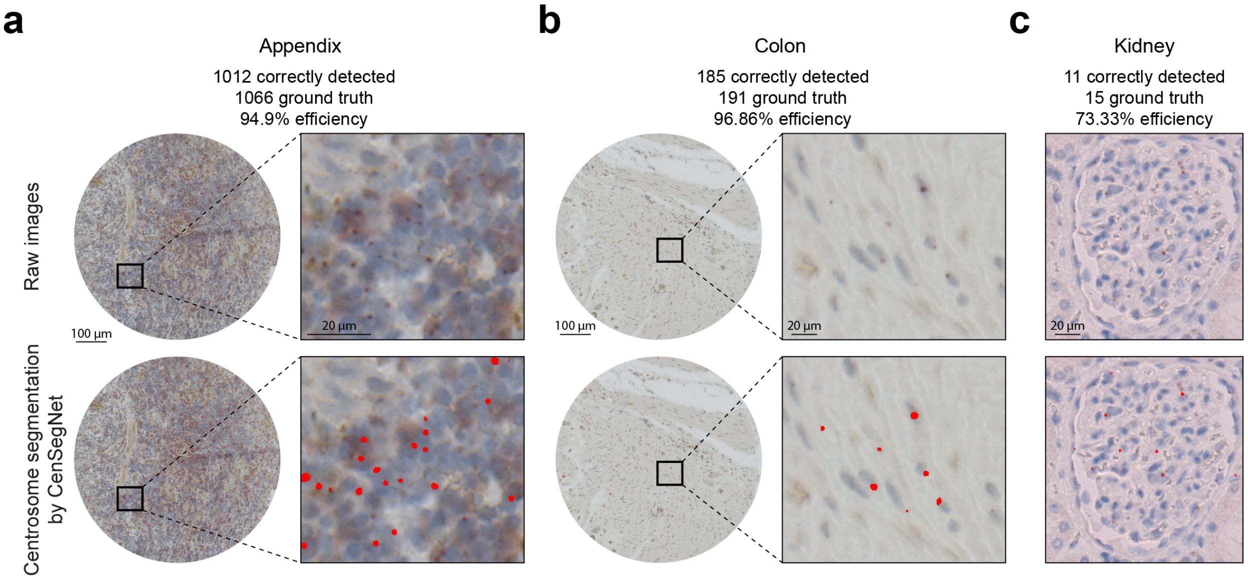


**Supplementary Fig. 17 CenSegNet performance in appendix, colon, and kidney tissues. a-c** Representative immunohistochemistry images showing CenSegNet centrosome segmentation performance in appendix (**a**), colon (**b**), and kidney (**c**) tissues stained for PCNT and counterstained with haematoxylin (nuclei). CenSegNet achieved detection efficiencies of 94.9% in appendix tissue [1,012 (correctly detected centrosome number)/1,066 (ground-truth centrosome number)], 96.9% in colon tissue [(185 (correctly detected centrosome number)/191(ground-truth centrosome number)], and 73.3% in kidney tissue [11(correctly detected centrosome number)/15(ground-truth centrosome number)]. Colon, kidney, and appendix tissue were included as orientation controls for breast tissue TMAs. Colon and kidney samples represent histologically normal tissue, confirmed by pathologist review.

#### Supplementary Fig. 18

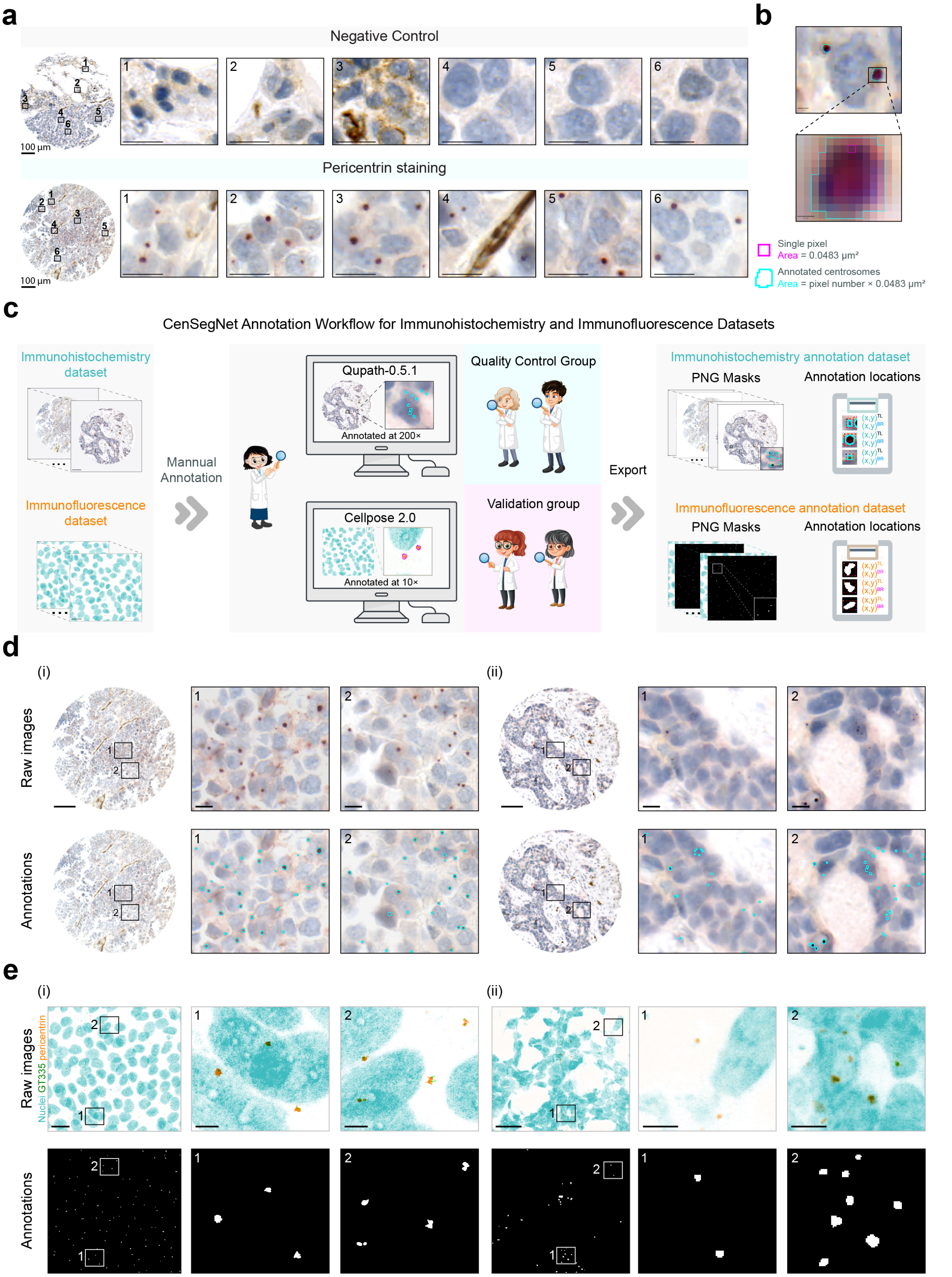


**Supplementary Fig. 18 Centrosome annotation across ground-truth datasets. a** Negative-control section processed in parallel without primary antibody, showing no specific staining. Nuclei are counterstained with haematoxylin (blue). Insets 1–6 show higher-magnification views of the boxed regions. **b** Representative section from the same patient stained for pericentrin.

Brown puncta indicate pericentrin-positive foci, frequently in a perinuclear distribution. Insets 1–6 show higher-magnification views of the boxed regions. Scale bars, 100 μm (main panels) and 10 μm (insets). **c** CenSegNet Annotation Workflow for Immunohistochemistry and Immunofluorescence Datasets. Centrosomes in immunohistochemistry dataset were annotated at 200× magnification, while those in immunofluorescence dataset were annotated at 10×. Annotations were then exported as PNG masks and accompanying text files specifying the location of each centrosome. TL indicates top left location; BR indicates bottom right location. Scale bars, 100 μm for immunohistochemistry dataset, 20 μm for immunofluorescence dataset. **d, e** Representative images with centrosome annotations in immunohistochemistry and immunofluorescence dataset. **d** (i) annotations of large centrosomes, (ii) annotations of small centrosomes. Scale bars, 100 μm for tissue cores, 10 μm for zoomed-in regions. **e** (i) annotations of human MECs, (ii) annotation of mouse mammary epithelium. Scale bars, 20 μm for raw images, 5 μm for zoomed-in regions.
